# Sequential roles for red blood cell binding proteins enable phased commitment to invasion for malaria parasites

**DOI:** 10.1101/2022.08.09.503398

**Authors:** Melissa N. Hart, Franziska Mohring, Sophia M. Donvito, James A. Thomas, Nicole Muller-Sienerth, Gavin J. Wright, Ellen Knuepfer, Helen R. Saibil, Robert W. Moon

## Abstract

Invasion of red blood cells (RBCs) by *Plasmodium* merozoites is critical to their continued survival within the host. Two major protein families, the Duffy binding-like proteins (DBPs/EBAs) and the reticulocyte binding like proteins (RBLs/RHs) have been studied extensively in *P. falciparum* and are hypothesized to have overlapping, but critical roles just prior to host cell entry. The zoonotic malaria parasite, *P. knowlesi*, has larger invasive merozoites and contains a smaller, less redundant, DBP and RBL repertoire than *P. falciparum*. One DBP (DBPα) and one RBL, normocyte binding protein Xa (NBPXa) are essential for invasion of human RBCs. Taking advantage of the unique biological features of *P. knowlesi* and iterative CRISPR-Cas9 genome editing, we determine the precise order of key invasion milestones and demonstrate distinct roles for each family. These distinct roles support a mechanism for phased commitment to invasion and can be targeted synergistically with invasion inhibitory antibodies.

## Introduction

Malaria threatens almost half the globe, with six species of *Plasmodium* parasites causing significant disease in humans (Ansari et al., 2016). Symptoms result from the blood stage of the parasite’s life cycle when motile merozoites invade red blood cells (RBCs), multiply within them, and then burst out (egress) to release new invasive merozoites. Since blocking invasion prevents parasite replication, a detailed understanding of how merozoites invade RBCs is essential to develop clinical interventions, such as vaccines and antibody therapies.

Video microscopy studies of *P. knowlesi (Pk)*, and later *P. falciparum (Pf)*, revealed merozoite invasion is a rapid process (∼60 seconds) with several morphologically-defined steps (Dvorak et al., 1975; Gilson and Crabb, 2009). First, the merozoite and RBC form a weak and reversible binding interaction, which quickly progresses into a stronger interaction causing the RBC to deform and ‘wrap’ around the merozoite. This may facilitate ‘re-orientation’ with realigning its apical end that houses specialised secretory organelles, micronemes and rhoptries, in perpendicular apposition to the RBC surface. Work in *Pf* identified a further step: fusion between the merozoite’s apex and the RBC before internalisation, visualised as a fluorescent signal at the parasite-host cell interface when RBCs are pre-loaded with a calcium-sensitive fluorescent dye (Weiss et al., 2015). Then, the merozoite actomyosin motor actively drives the parasite into the parasitophorous vacuole formed by invagination of the RBC membrane (Perrin et al., 2018). Invasion usually results in RBC echinocytosis, triggered by secretion of rhoptry contents or by perturbation of RBC homeostasis (Weiss et al., 2015). Within minutes the RBC returns to its biconcave shape, marking the end of a successful invasion (Dvorak et al., 1975; Gilson and Crabb, 2009).

The micronemes and rhoptries facilitate each step of invasion by sequentially releasing proteins that act as ligands for distinct receptors. Some of these ligands have been well characterised in *Pf*. For example, PfAMA-1 binds to PfRON2, its parasite-derived receptor inserted into the RBC plasma membrane, to form the moving junction, a molecular seal between merozoite and host cell that ‘travels’ backwards over the merozoite during internalisation (Lamarque et al., 2011; Srinivasan et al., 2011). Upon junction formation, the merozoite is presumed to be irreversibly committed to invasion.

Two key families of ligands predicted to mediate steps preceding junction formation are the reticulocyte binding-like (RBL/Rh) proteins and the Duffy binding proteins (DBP/EBA) (Tham et al., 2012; Weiss et al., 2015). *Pf* can express up to four DBP and five RBL orthologs. Simultaneously inhibiting PfRBL/DBP-receptor interactions prevents moving junction formation, RBC deformation and potentially merozoite re-orientation (Riglar et al., 2011; Tham et al., 2015; Weiss et al., 2015). However, significant redundancy within and possibly between the two families has hampered the dissection of their precise roles. Of all the PfRBL/DBPs, only PfRh5, an atypical RBL ligand restricted to *Plasmodium* species belonging to the subgenus Laverania appears to be essential for invasion, and likely performs a role distinct from the other RBL/DBPs at the point of merozoite-RBC fusion (Volz et al., 2016; Weiss et al., 2015). It remains unclear whether the other *Pf* RBL and DBP family members perform the same or distinct functions (Gao et al., 2013; Gunalan et al., 2020; Singh et al., 2010).

In contrast, evidence suggests that the RBL and DBP ligands of non-Laveranian Plasmodium *spp*, including the zoonotic parasite, *Pk* and closely related *P. vivax (*Pv), may have distinct functions. *Pk* expresses three DBP ligands (PkDBPα, PkDBPβ, and PkDBPγ), all highly similar in sequence to the single *Pv* DBP ligand, PvDBP. PkDBPβ and PkDBPγ are not required for invasion of human RBCs. However, PkDBPα and PvDBP are essential for invasion of human RBCs and bind to the Duffy antigen receptor for chemokines (DARC), limiting both species to replication in Duffy positive RBCs (Haynes et al., 1988; Miller et al., 1975; Mohring et al., 2019; Wertheimer and Barnwell, 1989). For *Pv*, RBL ligands interact with reticulocyte-specific receptors, such as PvRBP2b binding to TfR1 (Transferrin receptor 1), limiting this species to invasion of reticulocytes (Gruszczyk et al., 2018). *Pk* is not restricted to reticulocytes and can propagate in normocytes with one of its two RBL ligands, normocyte binding protein Xa (NBPXa) essential for growth in human RBCs (Moon et al., 2016). Thus, for both Pk and Pv, at least one RBL and one DBP are required for invasion of human RBCs. Early studies in *Pk* demonstrated that merozoites could deform Duffy negative cells, but not invade them, and that DBPα null merozoites cannot form a moving junction. However, it is unclear to what extent DBPα null merozoites can deform host cells, and which other steps of invasion are affected (Miller et al., 1975; Singh et al., 2005). Even less is known about the role of Pk/Pv RBL ligands, except that while NBPXa-null parasites can bind to the outside of human RBCs, they fail to invade them (Moon et al., 2016).

Recently we showed that merozoites can glide across RBC surfaces and that this active movement likely underpins RBC deformation (Yahata and Hart et al., 2021). However, the mechanisms by which gliding merozoites deform RBCs and transition into subsequent steps of invasion are unknown. Here we take advantage of the large size and clear polarity of *Pk* merozoites to explore the early steps of invasion and identify key milestones in host cell commitment. This provides a framework to investigate the roles of the PkRBL and PkDBP families using live microscopic analysis of fluorescently-tagged and conditional knockout (cKO) DBPα and NBPXa parasite lines, revealing these proteins have distinct locations once released from the micronemes. Finally, we demonstrate that NBPXa is essential for host cell deformation, while DBPα has a distinct function downstream of this, but ahead of rhoptry secretion and reorientation. These distinct roles provide a mechanism for staged commitment to host cell selection and invasion and identify these proteins as synergistic targets for therapeutic intervention.

## Results

### Deformation is associated with gliding motility and invasion success

We first used live-cell imaging of *Pk* merozoite-RBC interactions to delineate the morphological stages of invasion. From 20 egresses, 49% merozoite-RBC interactions (N = 134/275) displayed gliding motility across the RBC surface (Fig1A). Gliding was accompanied by RBC deformation in 67% of these interactions. On average this began within 2 seconds of gliding onset (median = 1 sec, 91 interactions) (Fig1A and S1A) and was characterized by a pinching or ‘wrapping’ of the host cell membrane across the width of the merozoite (Fig1B). This pinching appeared to originate at the merozoite apex and progressed towards the posterior (Video S1 from 2.0-6.5 sec). There were 5 unclear events (Fig1A), but we observed no clear instance of deformation without gliding, corroborating previous work showing gliding motility is required for deformation (Yahata and Hart et al., 2021). Importantly, all invasions (N = 40) were preceded by deformation, and as previously shown for *Pf* (Weiss et al., 2015), invasion success was positively correlated with deformation strength (Fig1B). These data suggest that both gliding and deformation are critical pre-cursors to *Pk* invasion.

**Fig1.**
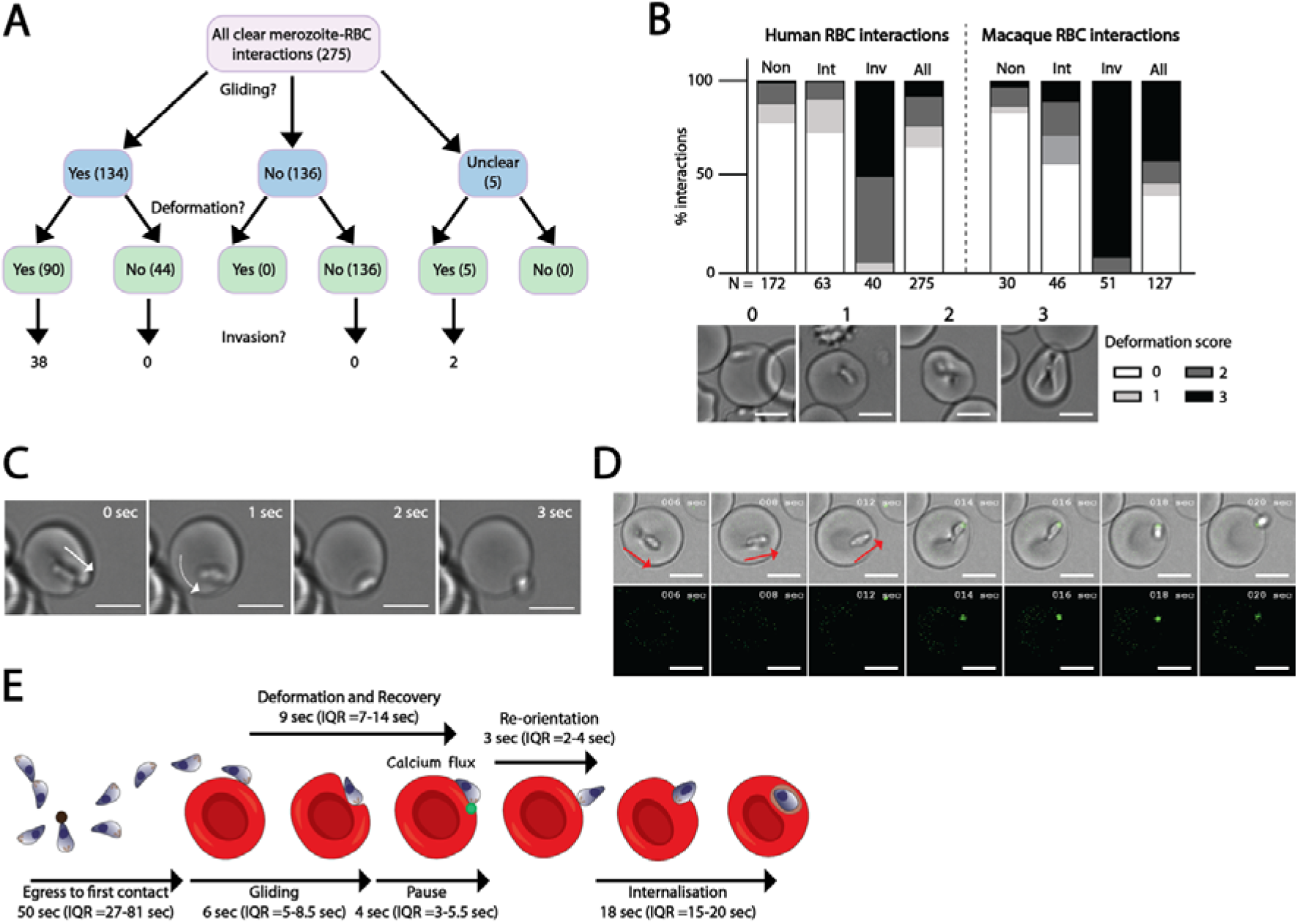
*P. knowlesi* as a model to delineate steps of RBC invasion. **(A)** Flow chart shows outcomes of all merozoite-human RBC interactions from 20 schizonts, 0-200 sec post egress. ‘Interaction’ = RBC contact lasting ≥ 2 sec. Gliding interaction = forward movement across RBC surface ≥ 5 sec. Events per category indicated in parentheses. **(B)** RBC deformation scores for human (left) or macaque (right) interactions based on extent of merozoite indentation/wrapping. 0 = no deformation; 1 = shallow indentation/membrane pinching; 2 = deeper indentation to the side of RBC/intermediate level of host cell membrane pinching around parasite; 3 = full host cell wrapping around merozoite. Bar chart indicates a breakdown of deformation scores for interactions from merozoites which never invade (non), intermediate interactions from merozoites that will invade on subsequent contacts (‘Int’), interactions leading directly to invasion (Inv), and all merozoite-RBC interaction (all). **(C)** Video S1 stills. Panel 1 shows RBC recovering from deformation and merozoite’s apical end firmly attached to the RBC membrane. White arrow tip indicates merozoite apex. Panels 2-4 show the merozoite pivoting on apical end (white curved arrow), until re-orientation is complete. **(D)** Video S3 stills. *Pk* merozoite displays a fluorescent signal between apex and Fluo-4-AM loaded RBC as gliding (red arrows = direction) comes to a pause (panel 4), but before re-orientation begins (panel 6). **(E)** Summary schematic depicting order and timings of each step of invasion. For all images, scale bars = 5μm.

While deformation strength is predictive of invasion success, it is unclear what factors determine a merozoite’s ability to initiate deformation. *Pk* merozoites frequently contact many human RBCs before commitment to invasion (Yahata and Hart et al., 2021). The deformation scores of interactions occurring prior to interactions resulting in invasion (‘intermediate interactions’ in Fig1B) are comparable to those for interactions made by merozoites which never invade (‘Non-invader events’). Fewer ‘intermediate interactions’ are observed when *Pk* merozoites invade macaque RBCs, which are highly permissive to *Pk* invasion, (Yahata and Hart et al., 2021) - suggesting that deformation strength is an indicator of host cell ‘suitability’ (for example, receptor availability or deformability), rather than merozoite ‘readiness’ to invade. A significant fraction of gliding interactions on human RBCs (44/134 events) did not progress to deformation (e.g., blue arrow in Video S2; Fig1A). Thus, whilst gliding is required for deformation, deformation of RBC membranes is not a prerequisite for traversal of RBC surfaces.

### Merozoite apex-RBC fusion precedes re-orientation

We examined individual invasion events to determine precisely when *Pk* merozoites are irreversibly attached to RBCs or ‘committed’ to invasion. *Pf* merozoites have been described to “pause” for several seconds on the host cell surface as deformation subsides but before internalisation begins (Gilson and Crabb 2009). During this stage, a merozoite-RBC fusion event takes place, visualised as a fluorescent signal appearing at the interface between the merozoite apex and Fluo-4-AM loaded RBCs (Weiss et al, 2015). However, the much smaller size and the more spherical morphology of *Pf* merozoites makes it difficult to determine when this step occurs relative to reorientation and junction formation.

Analysis of events immediately prior to internalisation showed that *Pk* merozoites appear to cease gliding motility – and thus forward movement across the host cell – as deformation subsides and a median 4 seconds (N = 16 events; S1B) prior to the RBC returning to its biconcave morphology. At this point, the stationary merozoite lies lengthwise across the RBC surface, with its apical end firmly pinned to the host cell surface (Fig1C, panel 1; Video S1 at ∼9 sec). The posterior end of the merozoite then dissociates from the RBC surface, while the merozoite pivots from the apical end until perpendicular to the RBC membrane (Video S1 from 9.5 – 11.5 sec and Fig1C, panels 2-4). Reorientation consistently began immediately after deformation ceased, and the RBC had returned to its original morphology (N = 28 events). Therefore deformation does not mechanically facilitate re-orientation as previously proposed (Dasgupta et al., 2014; Hillringhaus et al., 2019), but rather deformation and re-orientation are distinct, sequential steps.

Next, we examined merozoites invading Fluo-4-AM loaded RBCs. Strikingly, for all invasions where reorientation could be clearly visualised (N = 17/18 events from 12 egress events), a punctate fluorescent signal appeared at the apex of each merozoite ‘pinned’ to the RBC, a median 2 seconds before onset of reorientation (Fig1D; Video S3 at 14 sec; S1C). Frequently, we observed this punctate fluorescent signal move into the RBC with the invading parasite (e.g. Video S3 from 46 sec onwards), suggesting that this signal arises primarily from Fluo-4-AM travelling into merozoite compartments, such as the calcium-rich rhoptry lumen (Weiss et al., 2015). For 5/18 events, a faint but detectable fluorescent signal spread briefly to the RBC cytosol; however, this phenomenon was far less apparent than has been reported for *Pf* (Introini et al., 2018; Weiss et al., 2015). We observed no merozoite detachment after detection of Fluo-4-AM signal or reorientation. This indicates that commitment to invasion, and potentially also to moving junction formation, occurs before reorientation and thus earlier than suggested by data from *Pf* (Fig1E; Weiss et al., 2015; Introini et al., 2018).

### Propensity for strong deformation events underpin macaque RBC invasion success

We next investigated how *Pk* invasion and host cell selection differ between human and macaque RBCs. Whilst there was no significant difference in the length of time *Pk* merozoites spent invading either RBC (median = 33 sec vs 32 sec from first contact to completion of internalisation; p = 0.722; S1D), merozoites spent slightly longer deforming macaque vs human RBCs (median = 12.5 sec vs 9 sec; p = 0.016; S1E), and slightly less time actively entering macaque vs human cells (median internalisation = 15 sec vs 17.5 sec; p = 0.010; S1F).

Notably, a much higher proportion of interactions with macaque RBCs resulted in the strongest (score 3) deformation (N = 53/127 events for macaque cells versus 22/275 events for human)(Fig1B; Video S4). For both host cell types, strong deformation proved to be a reliable predictor of invasion success - with 89% of macaque (47/53 events) and 91% (20/22 events) of human RBC score 3 interactions progressing to invasion. This greater propensity for strong deformation suggests a greater proportion of macaque RBCs are amenable to invasion (e.g., RBC receptor availability, membrane tension etc.).

### Deletion of NBPXa prevents growth of *Pk* parasites in human but not macaque RBCs

Having established the morphological events leading to host cell entry, we sought to determine the role of NBPXa during invasion. Our previously nbpx*a* knockout line can only be maintained in macaque RBCs (Moon et al., 2016), so we established a conditional knockout (cKO) line that could be routinely maintained in human RBCs. A parasite line containing a dimerisable Cre recombinase (DiCre) was generated using CRISPR-Cas9 (Mohring et al., 2019) to insert a DiCre expression cassette into the *Pkp230p* locus (S2A & 2B). Adding rapamycin (Rap) to this line results in formation of active Cre recombinase, leading to excision of DNA sequence flanked by *loxP* sites. Subsequently, we iteratively modified this line to integrate a *loxP* sequence within the *nbpxa* intron and a HA tag and *loxP* site at the 3’ end of *nbpxa* (S2A & 2C; Fig2A).

**Fig2:**
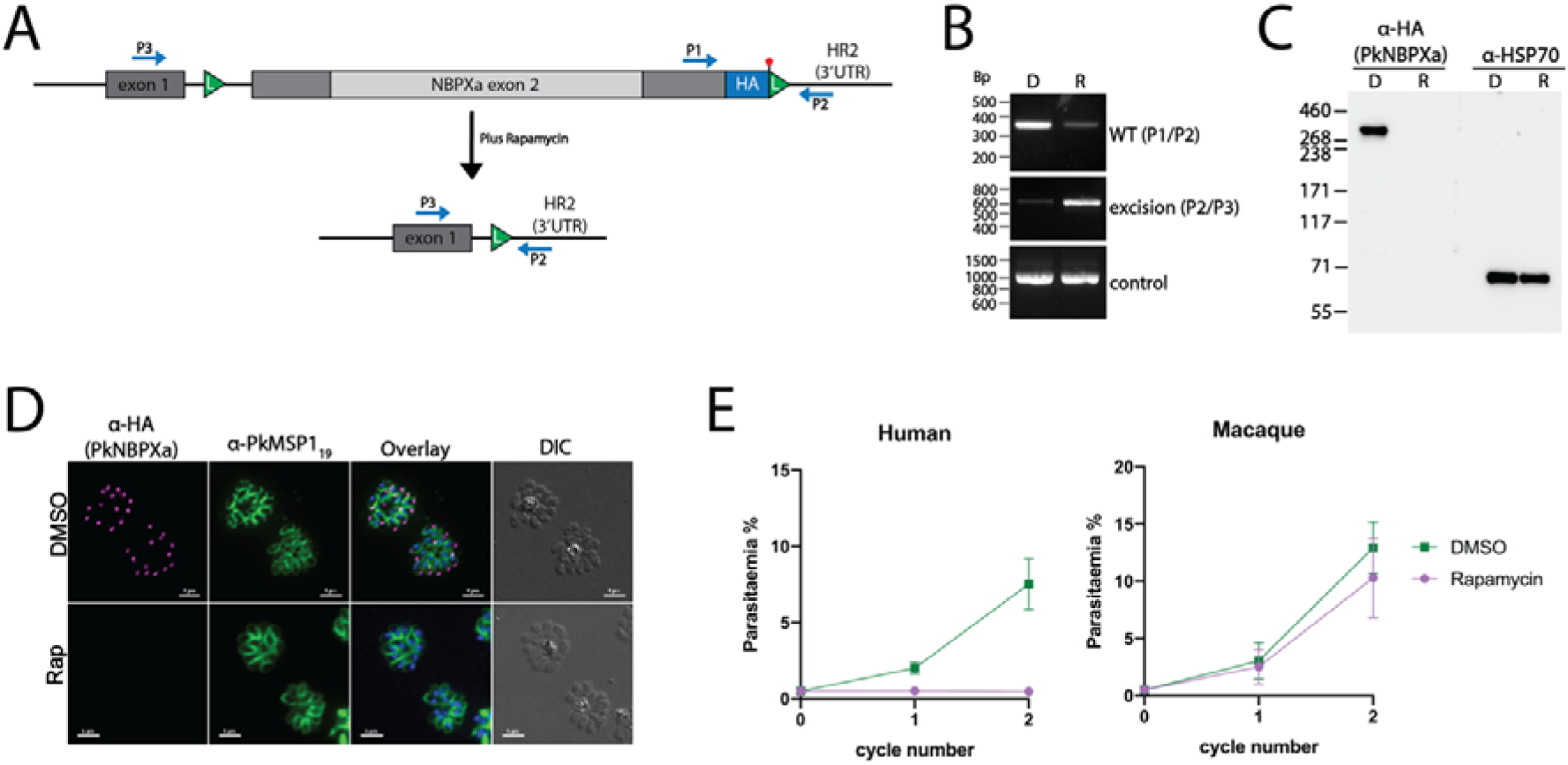
Efficient Rap-induced excision of floxed *NBPXa*. **(A)** Schematic depicting excision of 8.5kbp fragment between LoxP sites (L) upon Rap treatment of NBPXa cKO parasites. Diagnostic primer positions noted in blue (P1, P2, P3). **(B)** Diagnostic PCRs showing outcome of DMSO (D) or Rap (R) treatment of NBPXa cKO parasites. Primers P1/P2 identified non-excised parasites, P2/P3 detected successful excision and control bands amplify an unrelated locus. **(C)** Western blot showing loss of ∼325 kDa HA tagged NBPXa in Rap treated parasites. Anti-PfHSP70 used as a loading control. **(D)** IFAs showing ablation of NBPXa expression for Rap treated parasites. Parasites labelled with a rat anti-HA and rabbit anti-MSP1_19_ as a marker for mature, segmented schizonts. Scale bars = 5μm. **(E)** Growth of NBPXa cKO parasites in human (left) or macaque (right) RBCs measured at 24 and 48 hours after treatment with Rap or DMSO. Data shown are the mean of 5 independent experiments for human and 2 experiments for macaque cells. Error bars +/- SEM.

Synchronized ring-stage parasites were treated with 10nM Rap or carrier (0.005% DMSO) for 3 hours and samples were taken at the end of the first cycle (∼27-28 hours post-invasion) and analysed by PCR for successful *nbpxa* excision. Deletion of the floxed 8.4kb sequence removes most of *nbpxa* along with the C-terminal HA tag sequence, leaving only the signal peptide encoding exon 1 (Fig2A). PCR analysis demonstrated correct excision of the floxed *nbpxa* sequence in Rap-treated parasites (Fig2B). A fainter band was also detected in the control suggesting some formation of active Cre recombinase in absence of Rap. Full-length HA-tagged NBPXa (∼325 kDa) was detected by Western blot of mock-treated samples but was absent after Rap treatment (Fig2C). Immunofluorescence (IFA) analysis revealed (Fig2D) an overall excision rate of 93.5% (n= 22/342 HA positive cells for rap vs 258/271 DMSO).

Deletion of *nbpxa* led to a severe growth defect of *Pk* in human but not macaque RBCs (Fig2E). This is consistent with previous work and explained by the fact that NBPXb, the only nbpxa paralogue in *Pk*, binds macaque but not human RBCs, and thus complements the loss of NBPXa only for macaque RBC invasion (Moon et al., 2016). Rap treatment of wild-type parasites (S2D) revealed a minimal toxic effect of the drug, but this was not significant.

### NBPXa null merozoites are motile but fail to deform human RBCs

To understand why NBPXa null parasites fail to invade human RBCs, we examined merozoite-RBC interactions using live microscopy. These cKO merozoites were still capable of gliding across RBCs (Fig3A; Video S5): the median percentage of gliding merozoites per egress was 85% for both mock-treated and Rap-treated parasites (Fig3B; p= 0.5281). This result demonstrates that the merozoites were alive, with essential gliding and secretory systems intact, and that NBPXa is not required for the initial parasite-host cell interaction underpinning motility.

**Fig3.**
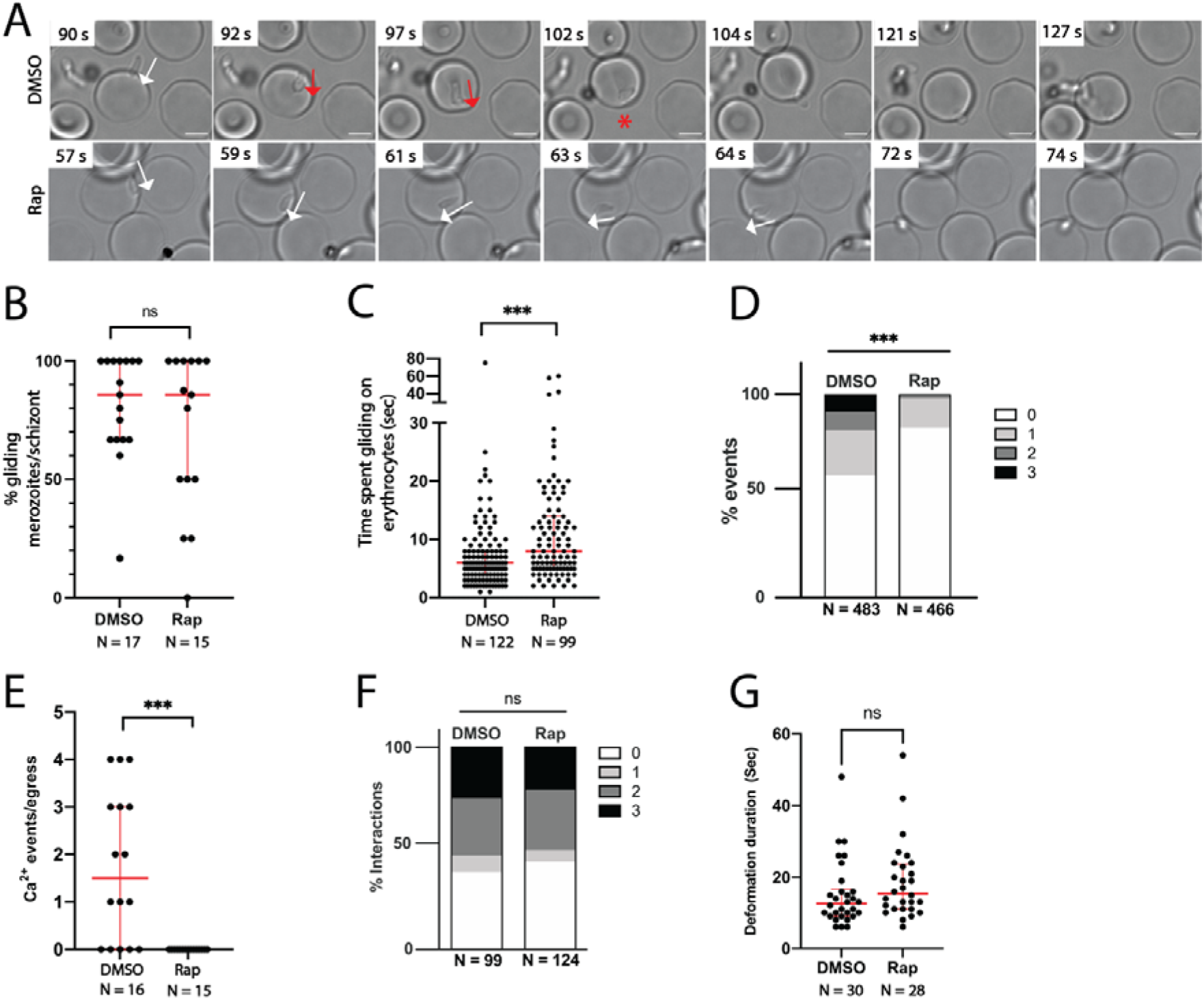
NBPXa is required for human RBC deformation. **(A)** Live stills from Video S5 showing DMSO vs. Rap treated NBPXa cKO merozoites interacting with human RBCs. White arrows indicate gliding motility without deformation. Red arrows indicate gliding motility with deformation. Red * = merozoite re-orientating for host cell entry. Interactions of DMSO vs. Rap treated NBPXa cKO merozoites with human RBCs were observed to compare **(B)** the % gliding merozoites per schizont (Mann-Whitney u-test) **(C)** the length of gliding interactions (Mann-Whitney u-test) **(D)** the proportion of strength 0-3 merozoite/RBC interaction events (chi-squared test) and **(E)** the number of Ca^2+^ events seen per egress when merozoites interact with Fluo-4-AM loaded RBCs (Mann-Whitney test). Invasion dynamics of the same lines with macaque RBCs were observed to compare **(F)** the proportion of strength 0-3 merozoite/RBC interaction events (chi-squared test) and **(G)** length of time merozoites spent deforming RBCs prior to internalisation (Mann-Whitney test). For all graphs, n indicated underneath. Thick red bars indicate the medians and thinner red bars indicate interquartile ranges. ns= non-significant, *** = p<0.0005.

On average, NBPXa null merozoites spent longer gliding on host cells (median = 8 sec) than mock-treated parasites (median = 6 sec; p = 0.0001; Fig3C) – a likely outcome as these merozoites could not invade. However, while gliding was unimpeded, they failed to deform host cells (Fig3A and 3D; Video S5). For mock-treated parasites, high scoring (score 2 and 3) deformations made up 17.6% of all interactions (N= 85/483 events). In contrast, no score 3 events were recorded for NBPXa-null parasites and score 2 interactions were less than 2% (N = 8/466 interactions) of all interactions. The proportion of score 1 deformations, which included very minor indentations, caused by collision of merozoites gliding into RBCs, was also significantly reduced (14.6% in NBPXa-null parasites versus 22.2% in mock-treated parasites; p= 0.0126). Therefore, NBPXa appears to be critical for strong deformation of human RBCs.

The lack of deformation correlated with a drastically lower invasion efficiency for NBPXa-null parasites. While 58 merozoites from 20 mock-treated schizonts invaded RBCs, only two invasions were observed for merozoites from 20 Rap-treated schizonts. Since the excision efficiency of the cKO line is about 94%, we conclude that these two invasive merozoites probably had intact NBPXa.

Aside from the two invasions, we also saw no evidence of parasite reorientation. We analysed the interaction of DMSO- and Rap-treated parasites with Fluo-4-AM loaded RBCs and while we observed an average of 1 to 2 Fluo-4-AM signal events per egress for mock-treated parasites (16 schizonts), none was seen for NBPXa-null parasites (15 schizonts; Fig3E). Therefore, we conclude that NBPXa is required for deformation of human RBCs and that merozoites lacking this ligand are unable to progress to downstream steps.

In contrast, NBPXa null parasites could deform and invade macaque RBCs as efficiently as controls. There was no significant difference in proportion of strong deformations or the length of time merozoites spent deforming host cells prior to invasion (Fig3F and 3G). These data support that NBPXb is able to compensate for the loss of NBPXa for macaque RBC invasion – strengthening the case that the invasion phenotype observed with human RBCs is directly due to NBPXa deletion, not malfunction of invasion machinery.

### DBPα functions downstream of NBPXa

To investigate the function of DBPα, we began by targeting its host cell receptor (DARC) with an antibody (anti-DARC; Fy6 epitope). Blocking this epitope inhibits both PkDBPα and PvDBP from binding to DARC and thus prevents RBC invasion (Mohring et al., 2019; Nichols et al., 1987; Smolarek et al., 2010). To visualise the effect of blocking DARC binding, we filmed merozoites interacting with human RBCs in the presence of either 5 μg/ml anti-DARC (IC50 = 0.3 μg/ml, S2E), or 5 μg/ml IgG isotype control.

Blocking DBPα-DARC interactions completely inhibited invasion (0 invasions, n = 20 egresses) compared to control parasites (56 invasions, n= 20 egresses). However, gliding and strong host cell deformation were unaffected by blocking DARC (Video S6; Fig4A and 4B). This was surprising, as given these merozoites were capable of strong deformation but could not invade, we might have expected them to perform a significantly greater number of strong deformations overall. However, only 3/15 merozoites which generated score 3 events performed an additional score 3 contact (vs. 1/17 for control). Notably, score 3 deformation lasted significantly longer for anti-DARC blocked (median = 18.5 sec) vs. control merozoites (median = 12 sec; p = 0.003; Fig4C), suggesting that if strong deformation is initiated but not resolved by reorientation and invasion, then the merozoite extends this process rather than moving on to another cell. Furthermore, anti-DARC blocked merozoites made fewer RBC contacts overall than NBPXa null parasites, which are not capable of performing strong deformation interactions (Fig4D). Thus, strong deformation and in particular, engagement of NBPXa with its host cell receptor, results in a ‘semi-committed state’ for the merozoite. In such cases, merozoites are unlikely to move to another cell and create a further score 3 interaction, which suggests that this prolonged strong deformation depletes resources required to support subsequent strong interactions (for example reduction of ATP or micronemal protein stores).

**Fig 4.**
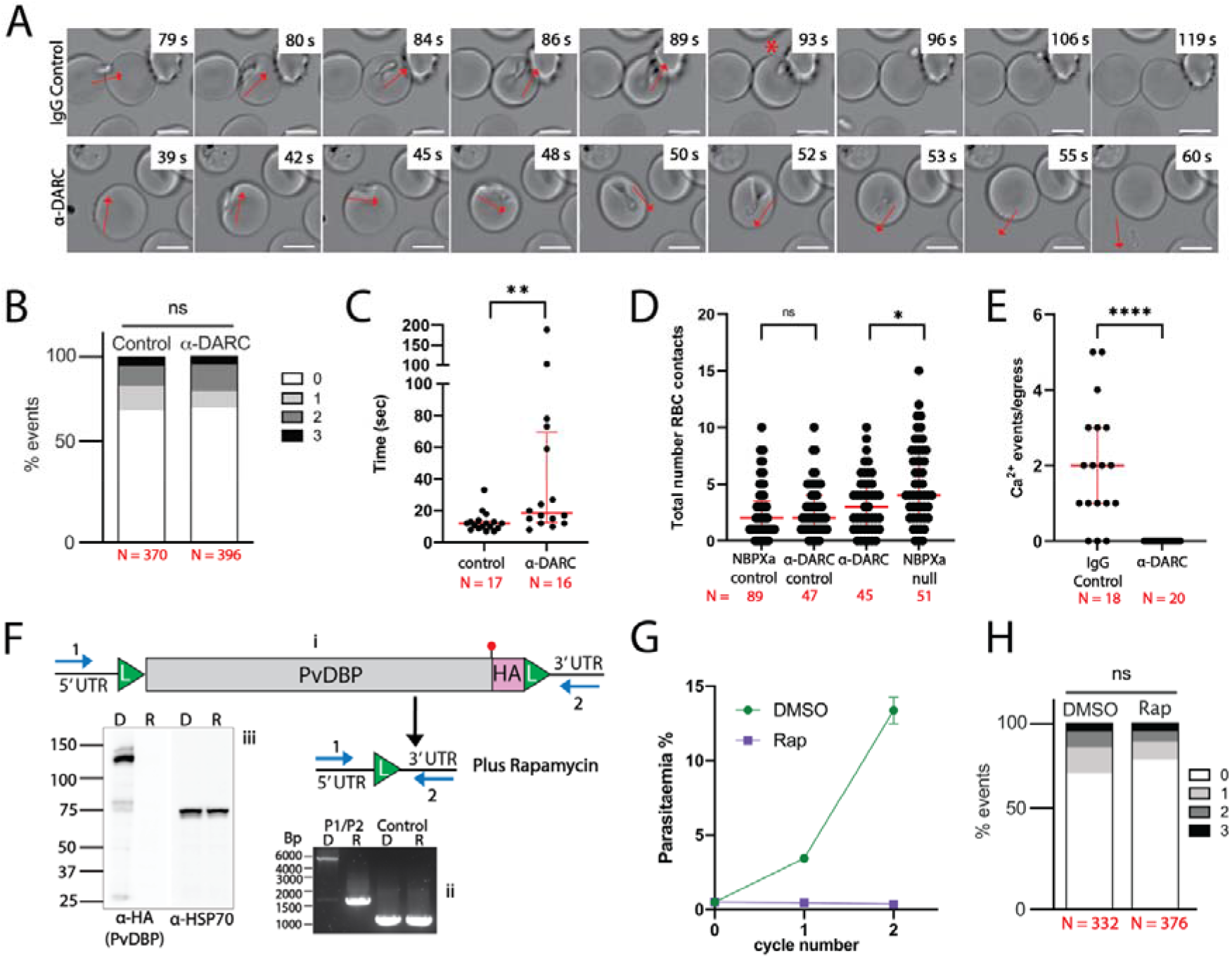
DBPα is required downstream of deformation. **(A)** Panels from Video S6 showing merozoite-RBC interactions in presence of 5 μg/ml IgG control (top) or anti-DARC (bottom). Red arrows indicate direction of gliding and deformation. Red * indicates onset of re-orientation and invasion. Invasion dynamics of merozoites with human RBCs in presence of IgG control or anti-DARC were quantified to compare **(B)** the proportion of strength 0-3 merozoite/RBC interaction events (chi-squared test) and **(C)** duration of each score 3 deformation event (Mann-Whitney). **(D)** Comparison of total number of RBC contacts made by NBPXa null vs anti-DARC blocked merozoites and controls over 2 min window post egress (Mann-Whitney U-test). **(E)** Number of Ca^2+^ events (per egress) seen when merozoites interact with Fluo-4-AM loaded RBCs with α-DARC vs IgG control (Mann-Whitney test). **(F)** Schematic (i) shows excision of *PvDBP* flanked by LoxP sites (L) upon Rap treatment of PvDBP cKO parasites. Diagnostic primer positions noted in blue (P1, P2) produce band shift of 4950 bp (non-excised parasites) to 1670 bp in PCRs (ii), with no change in unrelated control locus. Western blots (iii) probed with HA antibody detect ∼120 kDa PvDBP-HA in DMSO (D) but not Rap (R) treated parasites. Loading control = anti-PfHSP70. **(G)** Growth of PvDBP cKO parasites in human RBCs measured at 24 and 48 hours after treatment with either Rap or DMSO. Data mean of 3 independent experiments; error bars +/- SEM. (H) Comparison of strength 0-3 merozoite-RBC interaction events between Rap vs DMSO treated PvDBP cKO parasites (chi-squared test). For all graphs red bars indicated median + IQR, ns = non-significant, * = p<0.01, ** = p <0.005 and **** = p<0.0001.

DBPα null merozoites have been suggested to be able to reorientate on RBC surfaces (Singh et al., 2005). However, we observed no such events for anti-DARC blocked parasites, nor merozoite/RBC fusion events when fluo-4-AM loaded RBCs were treated with anti-DARC (Fig4E). In combination, these results suggest that DBPα acts downstream of NBPXa and is required for full commitment prior to re-orientation.

We next sought to generate a DBPα cKO line to compare the phenotype of DBPα null parasites with the effect of anti-DARC treatment. Initial attempts to flox *DBPα* failed due to inappropriate integration events associated with low complexity sequences; therefore, we replaced *PkDBPα* with a floxed *PvDBP* sequence, also expressing an HA tag (S2F), since *PvDBP* fully complements *PkDBPα* in human RBC invasion (Mohring et al., 2019). Rap treated PvDBP cKO parasites no longer expressed HA-tagged PvDBP (Fig4F) and could not proliferate in human RBCs (Fig4G). When examined by live microscopy, it was clear that PvDBP null merozoites could glide and deform host cells normally but could not progress beyond this step (Fig4H), like wild type parasites in the presence of anti-DARC.

### NBPXa and DBPα exhibit distinct localisations upon secretion from micronemes

Having established sequential roles for NBPXa and DBPα during invasion, we next sought to examine the localisation and secretion dynamics that underpin these roles. We tagged both NBPXa and DBPα at the C-terminus with the fluorescent marker, mNeonGreen (mNG) (Fig5A; S2A & S2G). We then added an HA tag to either NBPXa, AMA-1, or RON2 in the DBPα-mNG parasite line, and an HA tag to NBPXa in a previously established AMA-1mNG line (Yahata and Hart 2021; Fig5B; S2A, S2C & S2H) –using iterative CRISPR-Cas9 editing (Mohring et al., 2019).

**Fig5.**
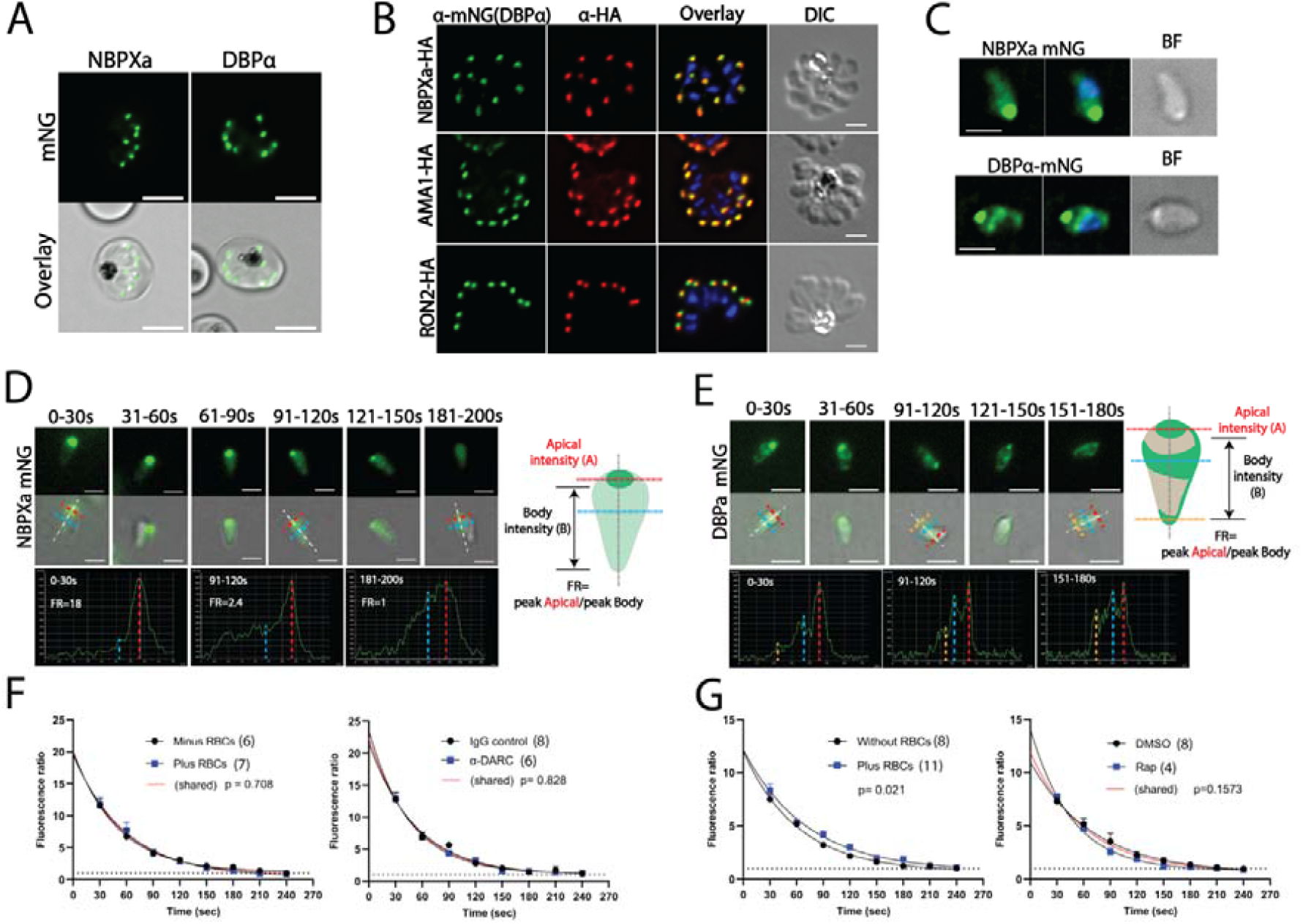
NBPXa and DBPα are secreted gradually and continuously post egress. **(A)** Image of mNeonGreen (mNG) tagged NBPXa and DBPα schizonts. Scale bars = 5μm. **(B)** Immunofluorescence assay showing HA tagged NBPXa, AMA-1, and RON2 (red), detected in schizonts using an anti-HA antibody. DBPα-mNG detected in same parasites (green) with an anti-mNG antibody. **(C)** Extended depth of focus images of NBPXa-mNG and DBPα-mNG in post egress merozoites. Live stills depicting NBPXa **(D)** or DBPα **(E)** secretion over a period of 3 minutes post egress. Secretion monitored by comparing the peak fluorescence intensity of the merozoite’s apical end, to the peak intensity at any point along its body (FR). Merozoite cartoons depict fluorescence pattern of each. NBPXa-mNG **(F)** or DBPα-mNG **(G)** FR values plotted over time. For both left-hand graph shows secretion +/- RBCs present. Right hand graph shows secretion with RBCs +/- anti-DARC antibody **(F)** or with RBCs and +/- NBPXa **(G)**. Measurements were recorded every 30 sec after egress (±5 sec either side) for all merozoites from a given schizont that were in focus at each timepoint. Each dot represents the average FR value/schizont. Error bars indicate ±SEM. FR values were fitted using non-linear regression; p values and number of schizonts (n) analysed for each group noted on graphs.

In schizonts both NBPXa and DBPα localised as a single fluorescent dot at the apical tip (wide end) of each merozoite (Fig5A and 5B). In dual-labelled IFAs DBPα appeared largely colocalised with both NBPXa (Pearson correlation R value = 0.94, n= 50 schizonts) and known micronemal marker AMA-1 (Pearson correlation R value = 0.96, n= 47 schizonts), confirming that both ligands are micronemal. Rhoptry neck marker RON2, overlapped slightly with a lower Pearson correlation R value, 0.84 (n=50 schizonts), demonstrating that it resides in separate organelles.

After merozoite egress NBPXa and DBPα had distinct locations (Fig5C). Over time, as NBPXa was released from the micronemes, the mNG signal gradually spread evenly across the merozoite (Fig5D), until the peak fluorescence intensity of the apical end normalised with that of the merozoite’s body (Fig5D, cartoon; Video S7). In contrast, DBPα exhibited a novel asymmetric surface localisation, accumulating at the widest circumference of the merozoite and the tip of its basal end, resulting in three discrete fluorescence peaks down the length of the merozoite (Fig5C and 5E). Interestingly, the localisation had distinct chirality with a solid stripe running down the length of one side of the merozoite (Fig5E). When DBPα-mNG tagged merozoites glide, this strip can be seen to rotate in and out of focus (Video S8). As secretion of DBPα progressed, the fluorescence ratio (FR) between the peak apical vs peak body intensity gradually normalised.

### Secretion of RBL/DBP proteins in *Pk* occurs independently of receptor engagement

Conflicting evidence in *Pf* suggests that initial receptor engagement by either RBLs or EBAs is required for secretion of the other RBC protein binding family (Singh et al., 2010; Gao et al., 2013). While in *Pf* RBL and DBP ligands are located in the rhoptries and micronemes, respectively, *Pk* and *Pv* RBL and DBP homologues are both located in the micronemes (Han et al., 2016; Meyer et al., 2009). Sequential release may still be possible in *Pk*, but would require segregation within subpopulations of micronemes. We observed NBPXa and DBPα secretion even without merozoites contacting RBCs (Videos S7 & 8). Therefore, engagement of NBPXa with its host cell receptor is not required for DBPα secretion, and vice versa. We next measured secretion rates of both NBPXa and DBPα under different conditions to explore impact of ligand-receptor binding. NBPXa secretion remained constant, regardless of the presence or absence of RBCs or when DBPα-DARC interactions were blocked with anti-DARC. For all conditions tested, peak apical vs peak body intensity normalised (FR approached 1) within ∼3-4 minutes after egress (Fig5F), corresponding to the window of maximum invasion following egress (S1G). Likewise, DBPα secretion progressed rapidly over a 3-4-minute window. A slight, but significant decrease in DBPα secretion speed was observed in the absence of RBCs. However, DBPα secretion remained constant when RBCs were present, but NBPXa was conditionally deleted (Fig5G). Therefore, our data demonstrate that micronemal secretion of RBL/DBPs is not dependent on receptor engagement for either NBPXa or DBPα in *Pk*.

### DBPα co-localises with the early moving junction before host cell entry

Several *Pf* DBL and RBL ligands localise to the moving junction of merozoites when treated with cytochalasin D (cyto D) to block internalisation, including PfEBA-175 and PfRh1 (Gunalan et al., 2020; O’Donnell et al., 2006; Triglia et al., 2009a). However, in our NBPXa mNG parasite line, the location of NBPXa did not change during merozoite internalisation. Apical NBPXa, likely representing un-secreted micronemal stores, moved into the RBC and did not form a junction-like structure (Video S9). Comparing the location of NBPXa-HA and AMA1-mNG by IFA, we found that cyto D-blocked merozoites had a single apical NBPXa signal, moving through the ‘double dot’ pattern seen from a cross section of AMA-1 labelled moving junction (Fig6A).

**Fig6.**
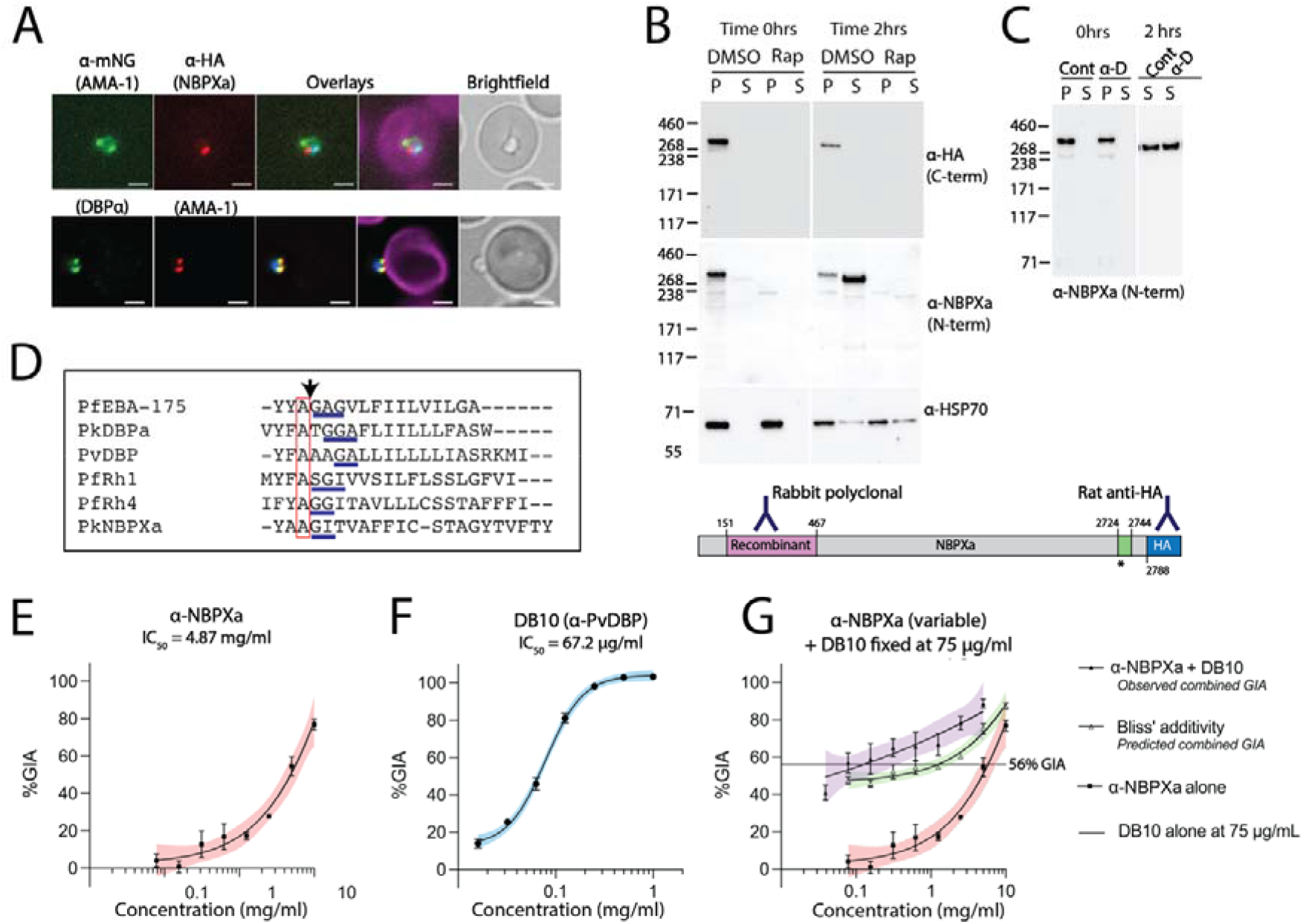
NBPXa acts prior to moving junction formation and can be synergistically targeted with DBP. **(A)** IFAs of *Pk* merozoites stalled mid-invasion with cyto D. Top panels depict AMA1-mNG + NBPXa-HA. Bottom panels depict DBPα-mNG + AMA1-HA. Overlays stained with wheatgerm agglutinin (Magenta). **(B)** Processed NBPXa is detected by western blot in culture supernatants (S) when egressing merozoites are allowed to re-invade fresh RBCs and **(C)** when invasion is blocked with anti-DARC (α-D). Cont = control IgG. For all blots, NBPXa was detected with rat anti-HA (targeting C-term) and/or rabbit anti-NBPXa raised against the NBPXa N-terminus (amino acids 151-467). Anti-PfHSP70 used as loading control. P = pellet fraction. Green region in schematic shows transmembrane domain, **\*** indicates putative ROM4 cleavage site. **(D)** Clustal alignment of several predicted RBL/DBP transmembrane domains. All contain a conserved alanine residue (red box), followed by putative helix destabilising motifs underlined in blue, which may serve as ROM4 recognition sites. All sequences apart from NBPXa from Baker et al., 2006 & O’Donnell et al., 2006. The experimentally determined PfEBA-175 cleavage position is indicated with black arrow. (O’Donnell et al., 2006). Parasite growth inhibition activity of anti-NBPXa **(E)** and anti-DB10 **(F)** was assessed individually and in combination using fixed DB10 concentration with increasing anti-NBPXa concentrations **(G)**. Results calculated from two independent experiments. Error bars indicate ±SEM. Coloured bands indicate 95% CI.

We tagged NBPXa at its C-terminus, which is proteolytically cleaved away from the ectodomain at some point during invasion. Attempts to localise the NBPXa ectodomain by IFA with a rabbit polyclonal antibody raised against the putative NBPXa RBC-binding domain (amino acids 151-467; Fig6B, schematic) were unsuccessful. However, western blots revealed that most of the NBPXa ectodomain was present in the culture supernatant as a single product following cleavage from its HA-tagged cytoplasmic tail (Fig6B). Consistent with imaging results, processing was unaffected by blocking invasion with anti-DARC antibody (Fig6C). This suggests that NBPXa is continuously secreted following egress and cleaved from the merozoite surface, potentially at the putative transmembrane ROM4 cleavage site, like its *Pf* counterparts (Baker et al., 2006; O’Donnell et al., 2006) (Fig6D).

DBPα-mNG largely co-localised with HA-tagged AMA-1 before host cell entry, with a circumferential ‘double dot’ structure at the merozoite-RBC interface characteristic of the early moving junction (Fig6A). However, live-cell imaging showed that the DBPα double dot pattern remained at the apex of invading parasites (See Video S10, 23 sec onwards) rather than travelling backwards along the parasite-RBC interface during entry - as is seen for typical moving junction markers, such as AMA-1. Thus, DBPα may play a role at or just before formation of the moving junction but is not part of the moving junction during entry.

### Simultaneous targeting of PkRBL and PkDBP invasion pathways inhibits invasion synergistically

Having established sequential roles for PkRBL/DBP family proteins during invasion, we sought to determine whether targeting both pathways would result in synergistic invasion inhibition. Rabbit polyclonal antibody targeting the putative RBC binding domain of NBPXa showed dose-dependent invasion inhibition of *Pk* with an IC_50_ of 4.9 mg/ml (Figure 6E), not dissimilar to inhibition by polyclonal anti-PvDBP RII antibodies (Mohring et al. 2019). To determine whether combining NBPXa and DBP antibodies would potentiate inhibition, we used our *Pk* PvDBP^OR^ line – combining our NBPXa polyclonal antibodies with DB10, a well characterised human monoclonal against PvDBP-RII (Rawlinson et al., 2019). Combining a fixed quantity of DB10 with a titration of NBPXa antibodies, we observed an increase in IC_50_ relative to the calculated Bliss Additivity curve – demonstrating clear synergistic activity between DBP and NBPXa antibodies (Fig6F & 6G).

## Discussion

The tractability of *Pk* for genetic manipulation, together with the relatively large size of its merozoites have enabled us to delineate the morphological and molecular processes underlying invasion. Our results show that gliding motility, deformation, merozoite-RBC fusion event, and re-orientation, are distinct, essential steps leading to host cell entry. We have also shown for the first time that although the *Pk* RBL and DBP ligands may be secreted simultaneously, following secretion and during invasion they exhibit distinct localisations. Our cKO data also demonstrates NBPXa and DBPα have distinct roles, with DBPα functioning downstream of NBPXa.

NBPXa null merozoites retain their ability to glide but cannot deform human RBCs, demonstrating that both gliding motility and NBPXa-receptor interactions are required for deformation. Yet how does NBPXa mediate deformation? NBPXa may be coupled to the merozoite’s actomyosin motor, as suggested but not confirmed for the *Pf* RBL/DBPs (Diaz et al., 2016; Pal-Bhowmick et al., 2012). Alternatively, NBPXa-receptor binding, in combination with the rotational movement of gliding motility (Yahata and Hart et al., 2021), may pull the RBC membrane around the merozoite. Enzymatic cleavage of NBPXa at its putative rhomboid ROM4 site may sever these links, allowing the merozoite to continue its forward trajectory. ROM4 processing of *Pf* RBL and DBP ligands is predicted to occur during host cell entry, to shed the ectodomains of these proteins during internalisation (Baker et al., 2006; Favuzza et al., 2020; O’Donnell et al., 2006; Triglia et al., 2009b). However, cleavage of these ligands occurs whether or not *Pf* merozoites re-invade (Favuzza et al., 2020), in line with what we observe for NBPXa. Thus, rhomboid cleavage of RBL ligands may primarily facilitate deformation instead, and the role of the RBLs may be to increase the “stickiness” of the parasite to create torsion-driven wrapping.

We also demonstrate that blocking DBPα-DARC interactions prevents merozoite apical fusion with the host cell and reorientation on the RBC surface. Interestingly, this mimics the consequence of disrupting the *Pf* Rh5/CyRPA/Ripr complex (Weiss et al., 2015; Volz et al., 2016), though *Pk* lacks an PfRh5 ortholog. However, *Pk* does have orthologs of PfCyRPA and PfRipr, which are two essential proteins (Chen et al., 2011; Knuepfer et al., 2019; Reddy et al., 2015; Volz et al., 2016), Further work will be required to dissect the molecular steps leading to merozoite reorientation and to determine the function of DBPα relative to that of the PkRipr-complex.

The implications for understanding *Pf* invasion are more complex, with previous evidence suggesting overlapping functions for that DBP/EBA and RBL/RH ligands. Improved deformation may enable a lower affinity EBA interaction to be successful and vice versa. It is also plausible that *Pf* uses these families in a more interchangeable way, underpinned by some key differences such as duplicated RBC-binding domains in EBA175 (vs single domain in DBPs), the distinct location and secretion timings of the PfRH and PfEBA proteins (Cowman et al., 2017). Nevertheless, visualising the key steps of invasion which are difficult to display in *Pf* provides a new perspective on key events shared across the genus - most notably that deformation and reorientation are clear and separate events, and that a merozoite-host cell fusion event and potentially tight junction formation occur before reorientation. Cross-species comparisons of invasion provide an invaluable tool to understand the biology underpinning these complex interactions and how to target them across all *Plasmodium* species.

Finally, our results reveal a mechanism for a stepwise commitment to invasion, which may underpin *Plasmodium* species host cell tropism (Figure 7). For all species, RBL and DBP repertoires determine the range of RBCs amenable to invasion (Tham et al., 2012). Most *Pf* RBL/DBP receptors are broadly distributed across human RBCs of different age and blood type, enabling this species to infect a wide range of individuals (Tham et al., 2012). However, some non-Laveranian species are restricted to growth in RBC subsets including *Pv* and *P. ovale*, to reticulocytes (Collins and Jeffery, 2005) and *Pk* and *Pv*, in most instances, to Duffy positive RBCs.

**Fig7.**
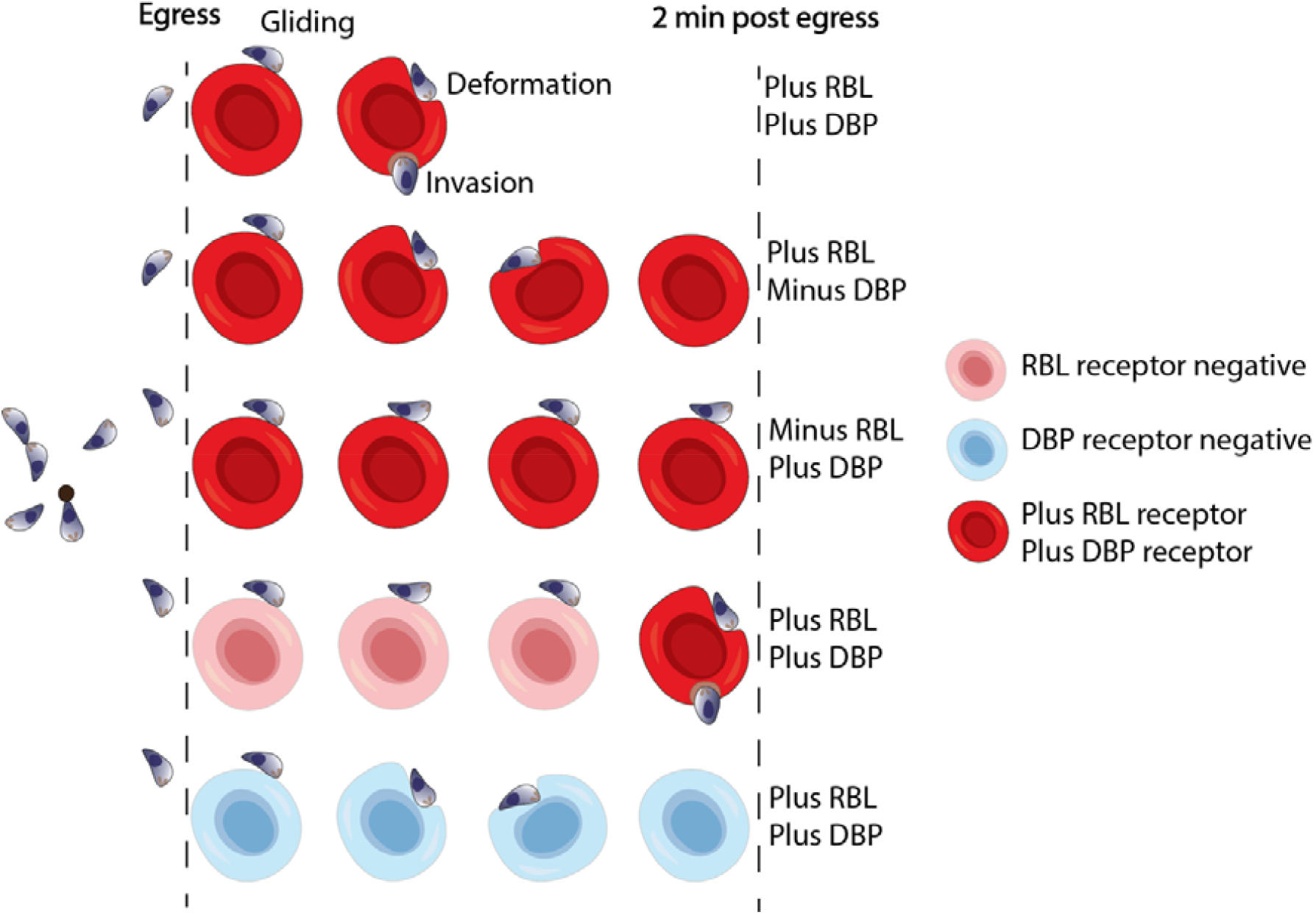
A model of how RBL and DBP ligands mediate phased commitment to invasion. Both in the absence of NBPXa and when host cells lack appropriate RBL receptor(s), *Pk* merozoites contact a greater number of RBCs overall, increasing their chances of interacting with a suitable RBC. In contrast, DBP null merozoites, or WT parasites contacting Duffy negative host cells, form semi-committed interactions, thereby reducing their chances of contacting a suitable host cell.

Yet, how do merozoites ensure they detect, bind to, and invade rare RBC subsets without depleting valuable resources attempting to invade unsuitable host cells? Our data indicate that NBPXa, and by extension, essential RBLs of other non-Laveranian species, mediate the first phase of commitment to invasion: host cell deformation. NBPXa null *Pk* merozoites do not deform RBCs on contact, and subsequently make a greater number of RBC contacts overall. In contrast, when NBPXa is present but DBP-receptor interactions are blocked, merozoites spend more time interacting with RBCs they deform but cannot invade, reducing their chances of subsequent successful interactions (Fig7). *In vivo*, this property of RBLs may present the parasite with a distinct advantage: the ability to sample host cells efficiently by quickly gliding over RBCs lacking an appropriate RBL receptor and thereby saving finite resources for invasion of an appropriate receptor-positive cell. This may explain how *Pv* merozoites can efficiently invade reticulocytes (which form ∼1% of RBCs in peripheral blood), as the reticulocyte-specific RBL proteins ensure the merozoite is “blind” to normocytes. Testing this hypothesis will be challenging due to our inability to culture *Pv in vitro* but combining orthologous gene replacement approaches in *Pk* and *P. cynomolgi* and using *Pv* in *ex vivo* experiments, could offer routes to address this.

The proposed sequential, phased commitment to invasion presents key opportunities for targeted vaccine design. In recent years, work to target *Pf* RBL and DBP proteins (apart from PfRh5)as vaccine candidates has been hampered, by both redundancy and polymorphism (Beeson et al., 2016; Duffy and Patrick Gorres, 2020). Our work shows that this is not the case for *Pk*. The growth inhibition assays with anti-PvDBP and anti-PkNBPXa antibodies demonstrated that blocking two critical, sequential invasion steps in combination dramatically curbs parasite growth compared to targeting either ligand individually. These results may be translatable to other *Plasmodium* species. Targeting a critical PvRBL ligand (Chan et al., 2020) together with PvDBP may be the key to developing an effective *Pv* vaccine, building upon promising results from recent PvDBP Phase II clinical trials (Hou et al., 2022).

## Acknowledgements

MNH was supported by a Bloomsbury Colleges Studentship. FM and RWM were supported by the UK Medical Research Council (MRC Career Development Award to RWM MR/M021157/1). JAT was supported by a Wellcome Trust Sir Henry Wellcome Fellowship. GJW and NM-S were funded by the Wellcome Trust (grant 206194). SMD was supported by a UK Medical Research Council LID studentship. HS was supported by UK Medical Research council Project Grant MR/P010288/1. The authors would like to thank Anthony A. Holder for critical reading of this manuscript.

## Author contributions

MNH, RWM, and HRS conceived and designed research. MNH, FM, and SMD performed research. MNH RWM, JAT, FM and EK contributed to conceptualisation and methodology. NM and GJW contributed the generation of recombinant NBPXa for antibody production. MNH and RWM wrote the paper, and all authors contributed to the manuscript and analysed the data.

## Methods

### Recombinant NBPXa production

A gene fragment of PkNBPXa encoding amino acids 151 to 467 (IDRILD… to …DALKDK) were flanked by unique NotI and AscI restriction enzyme sites and cloned into a pTT3-based mammalian expression plasmid between a mouse antibody N-terminal signal peptide and a C-terminal tag that included a protein sequence that could be enzymatically biotinylated by the BirA biotin ligase and 6-His tag for purification [PMID: 24620899]. The ectodomains were expressed as soluble recombinant proteins in HEK293 cells as previously described (Wanaguru et al., 2013). To prepare purified proteins for immunization spent culture medium containing the secreted ectodomain was collected from transfected cells, filtered and purified by Ni^2+^-NTA chromatography using HisTRAP columns using an AKTAPure instrument (GEHealthcare). Proteins were eluted in 400 mM imidazole as previously described [PMID: 31952523], and extensively dialysed into HEPES-buffered saline (HBS) before being quantified by spectrophotometry at 280 nm. Protein purity was determined by resolving purified protein by SDS–PAGE using NuPAGE 4–12% Bis Tris precast gels (ThermoFisher) at 200 V. Purified proteins were aliquoted and stored frozen at −20□°C until used. Where enzymatically monobiotinylated proteins were required to determine antibody titres by ELISA, proteins were co-transfected with a secreted version of the protein biotin ligase (BirA) as previously described [PMID: 27226583], and extensively dialysed against HBS and their level of expression determined by ELISA using a mouse monoclonal anti-His antibody (His-Tag monoclonal antibody, 70796, EMD Millipore) as primary antibody and a goat anti-mouse alkaline phosphatase-conjugated secondary (A3562, Sigma-Aldrich).

### NBPXa antibody generation and purification

Rabbit anti-PkNBPXa antibodies were raised (Covalab) against the recombinant protein described above and subsequently purified with an immunoaffinity column prepared by coupling recombinant protein to 1ml of activated sepharose beads. In brief, anti-NBPXa antibodies were purified by diluting equal parts rabbit serum and 10mM Tris (pH 7.5) and running over column 3 times. The column was then washed by addition of 20mL 10mM Tris (pH 7.5), then again with 20mL 10mM Tris (pH 7.5) with 0.5mM NaCl. Bound anti-PkNBPXa was eluted by passing 10mL glycine over the column and collecting in a tube containing 700μL 1M Tris HCl (pH 8.0) to neutralise the glycine. Eluted protein was concentrated and buffer exchanged into incomplete media (ICM) using the Amicon® Pro Purification System with a 50,000 molecular weight cut-off (MWCO), as per the manufacturer’s instructions. Final antibody concentration was read using the DeNovix DS-11 Series microvolume spectrophotometer, blanked with ICM and with the antibody mass extinction coefficient (E1%) set to 14.4 for rabbit IgG. Concentrated, buffer exchanged IgG was stored at -20°C until use.

### Generation of DNA constructs for transfection

#### Cas9 guide plasmids

All Cas9 guide RNA sequences (NBPXa N-term: AAATTCATGAACCCCAATTA; RON2 C-term: ACGCCCGCATACAGATGTA; and DBPa-C-term: CATGCAGCAGTTCACCCCCC), apart from one targeting the NBPXa C-terminus (GTAACGAATATATATGAGTA), were inserted into vector pCas9/sg according to previously published methods (Mohring et al., 2019). Because of difficulty cloning the NBPXa C-term guide into pCas9/sg, this sequence was inserted into a second Cas9 vector, pCas9HF/sg. This vector contains all component of pCas9/sg but has a smaller backbone and a high fidelity (HF) Cas9 sequence instead of the wild type version. Insertion of the NBPXa C-term guide into pCas9HF/sg was achieved by amplifying the entire guide cassette using overlapping primers (Supplementary Tables 1 and 2) containing the guide sequence. This PCR product was digested with PspXI and BstEII (NEB) and ligated into the pCas9HF/sg using T4 DNA ligase (Promega). Previously published guide plasmids pCas9_p230p (Mohring et al., 2019) and pCas9_AMA1_Cterm (Yahata and Hart et al., 2021) were used to target the *p230p* and *AMA-1* loci, respectively.

#### Donor DNA constructs

##### Generation of DiCre cassette Donor DNA

The donor plasmid for introducing the DiCre cassette into *Pkp230p* was generated in multiple steps. Primers and templates used to amplify each component are listed in Supplementary tables 1 and 2. First, 5’ and 3’ 400 bp homology regions flanking the *Pkp230p* guide were amplified from Pk A1-H.1 genomic DNA and inserted into a plasmid backbone containing a multiple cloning site (synthesized by Geneart, Thermofisher) via SacII/SpeI and BlpI/NcoI restriction sites, respectively.

To generate a Cre60 cassette, the *PkHSP70* 5’UTR sequence was amplified and inserted into a pGem-T Easy vector (Promega) followed by the *Cre60* sequence via restriction cloning between XhoI/KpnI sites, and the *PfPbDT* 3’UTR sequence between KpnI/BlpI sites. Likewise, to generate a *Cre59* cassette, the *PkEF1α* 5’UTR sequence was amplified and inserted into the pGem, followed by the *Cre59* sequence (XhoI/KpnI) and *PfHRP2* 3’UTR sequence (KpnI/SpeI). Finally, the PkHSP70 5’UTR-Cre60-PbDT 3’UTR cassette was introduced between *p230p* homology arms via SpeI/BlpI restriction sites, and the PkEF1α 5’UTR-Cre59-PfHRP2 3’UTR cassette via SpeI/NotI restriction sites.

##### Generation of *PkNBPXa, PkDBPα, PkRON2, PkAMA-1*, and *PvDBP* tagging and cKO donor DNA constructs

Donor DNA templates used to modify *PkDBPα, PkNBPXa, PkRON2, PkAMA-1*, and *PvDBP* with either tags or LoxP sites were generated by overlapping PCR, according to methods outlined in Mohring et al., (2019) and using primers and templates listed in Supplementary Tables 1 and 2. The NBPXa-mNG, NBPXa-HA, RON2-HA, and AMA1-HA constructs were transfected as PCR products. Prior to transfection, the DBPα-mNG, NBPXa-NtermLoxP, and NBPXa-CtermLoxP/HA constructs were first cloned into plasmid backbone (Geneart, Thermofisher Scientific), containing a multi-cloning site with SacII, NotI, and EcoR1 restriction sites. The final PCR amplification step of the DBPα-mNG construct added a SacII restriction site to the 5’ end of the construct and an EcoR1 site to its 3’ end for insertion into vector backbone. Likewise, the final PCR amplification step of NBPXa-NtermLoxP and NBPXa-CtermLoxP/HA constructs added a SacII restriction site to the 5’ ends and a NotI restriction site to the 3’ ends of both constructs for introduction into PkB1 by restriction cloning.

In order to generate the PvDBP cKO donor template, plasmid pDonor_PvDBP^OR^ (Rawlinson et al., 2019) was sequentially modified to introduce LoxP sites flanking the *PvDBP* sequence. The plasmid was first linearized with BstBI and NotI to remove 90bp at the C-term end of PvDBP^OR^. This 90 bp was replaced with an insert including a HA tag-LoxP sequence. A second LoxP sequence was introduced before the *PvDBP* start codon by removing homology region 1 (HR1) via SacII/SpeI and replacing it with a HR1-LoxP sequence.

#### In vitro maintenance and synchronization of *P. knowlesi* parasites

Blood stage A1-H.1 P. knowlesi parasites were cultured in human erythrocytes (UK National Blood Transfusion Service) at a 2% haematocrit with custom made RPMI-1640 medium, supplemented with 10% Horse Serum (v/v) and 0.292 grams/litre (2 mM) L-glutamine according to previously established methods (Moon et al., 2013). Parasites were tightly synchronized for experiments by first purifying schizonts with a density gradient centrifugation step using 55% Nycodenz (Axis-Shield) and allowing parasites to re-invade fresh RBCs over a 1-2 hr window. Next, newly formed ring stages were isolated by incubation with 140mM guanidine hydrochloride for 10 min to kill all parasites older than 5 hrs old, according to a protocol outlined by Ngernna et al., (2019)

### Transfection and genotyping of *P. knowlesi* parasites

Transfections were performed with 10-20 μl mature schizonts, 10 μg of the Cas9 plasmid, and 20 μg of the donor DNA using the Amaxa 4-D Nucleofactor X machine (Pulse code FP158) and P3 Primary Cell Kit (Lonza), according to protocols described by Moon et al. (2013) and Mohring et al. (2019). Transfected parasites were cultured with 100 nM pyrimethamine for 5 days following transfection in order to select for parasites that had taken up the Cas9 plasmid. A minimum of 1 week later, cultures were subsequently treated with 1 μM 5-Fluorocytosine for 7 days to eliminate parasites still harbouring the Cas9 plasmid (Mohring et al., 2019). Finally, transgenic parasites were cloned by limiting dilution (Moon et al., 2013).

Transfected parasites were analyzed by diagnostic PCR using GoTaq Green master mix (Promega) and the following conditions: 3 min at 96°C, then 30 cycles of 25 s at 96°C, 25 s at 55°C, and 1 min/kb at 64□C. Diagnostic primers along with expected PCR product sizes are listed in Supplementary Tables 1 and 2.

### Rapamycin treatment of conditional knockout lines

Tightly synchronised ring stage *P. knowlesi* cultures were split in two and incubated in complete media containing 10 nM rapamycin (Sigma-Aldrich) or the equivalent volume of its carrier, DMSO (0.005%). Parasites were treated for a minimum of 3 hours at 37°C, before drug or DMSO containing media was removed and cultures were re-suspended in fresh, drug-free complete media.

Diagnostic PCRs were carried out using GoTaq Green to detect excised or non-excised parasites after drug or mock treatment. For PkNBPXa cKO parasites, primers Fwd-GTTTCACTATTGAAGGATAATTTTAGGAAAGG-3’ and rev-GTATTACGGTTAATATGTTTTAACGTAACTGG were used to detect excised parasites, giving an expected product size of 606bp. Primer pair Fwd-ACTGCCGGATACACAGTCTTTAC and Rev-GTATTACGGTTAATATGTTTTAACGTAACTGG were used to detect non-excised parasites, with an expected product size of 404 bp. For PvDBP cKO parasites, primers Fwd-CCATGTACACGATTTGTGTACTTATAGAATC and Rev-GTAGGGAACATTTCTTTCTGCGG were used to detect both WT (expected product size = 4950 bp) and excised (expected product size = 1670 bp) parasites.

A positive control using primers Fwd-CCCGGGGCGTTTTCGCGTATCTGCGCTTTTTC and Rev-CCTAGGGGACAATATATCCTCACAGAACAACTTG, which amplified a 1043bp product from the PkMTIP locus, was also included in each set of reactions.

### Parasite multiplication assays

To determine the multiplication rate of mutant parasites, ring stage DMSO and rapamycin-treated cultures were adjusted to a 0.5% parasitaemia and 2% haematocrit and were grown in triplicate in 96 well plates in a gassed chamber at 37□C. After 24 hours, a starting sample was taken and stained with SYBR Green I (Life Technologies) before measuring parasitaemia by flow cytometry (FACS). A second sample was taken 26 hours later, and a final sample was taken roughly 26 hours later again. Assays were analysed on an Attune NxT flow cytometer using FACSDiva 6.1.3 software.

To determine the invasion inhibitory potential of anti-DARC (2C3 clone, Absolute Antibody), anti-DB10 (Rawlinson et al., 2019) and anti-NBPXa antibodies, growth inhibition assays (GIA) were carried out using an LDH assay described in Mohring et al., 2020. Bliss’s additivity was calculated using equation described in Williams et al., (2012).

### Immunofluorescence assay (IFA) analysis

Late stage schizonts were thinly smeared on glass slides, air dried, and fixed in 4% paraformaldehyde (PFA) in PBS for 30 minutes, followed by 0.1% Triton X-100 in PBS for 10 min. Slides were subsequently blocked in 3% BSA in PBS overnight. All primary and secondary antibodies were diluted in blocking solution and incubated sequentially for 1 hr at room temperature followed by three washes in PBS for 10 min each. Slides were incubated with the following antibodies: mouse α-mNeonGreen (1/500; 32F6, Chromotek) followed by α-mouse IgG Alexa Fluor 594, rat α-HA (1/500; 3F10, Sigma), followed by α-rat IgG IRDye 680 RD (LI-COR Odyssey), and rabbit α-rabbit PkMSP1 (1/2000; Ellen Knuepfer, Francis Crick), followed by α-rabbit IgG Alexa Fluor-488. After a final wash in PBS (3x 10 min), slides were mounted in ProLong Gold Antifade Mountant containing DAPI (Thermo Fisher Scientific).

For in solution IFAs, merozoites were captured mid-invasion by treating egressing cultures with 100 nM cytochalasin D (Sigma) for 30 min and then gently spinning cultures down at 1500 x g for 1 minute. Parasite pellets were re-suspended in a 4% PFA/0.0025% glutaraldehyde solution in PBS, and then loaded into a poly-L-lysine coated μ-Slide VI 0.4 (Ibidi). Parasites were incubated in fixative solution for 30 min before fixative was removed by pipetting the solution out of the channel slides. Channels were then washed 5x by gently pipetting PBS into the channel and then removing it in the same manner as the fixative. Subsequently, parasites were incubated with 0.1% Triton X-100 in PBS for 10 min, followed by 5 wash steps in PBS. Washed parasites were blocked with 3% BSA in PBS for at least 1 h. Samples were subsequently incubated with primary and secondary antibodies as described above, with 5 x wash steps in between each incubation. After the final wash steps, parasites were incubated in PBS containing Hoechst 33342 (Cell Signaling Technology) for half an hour prior to imaging. All samples were imaged with a Nikon Ti E inverted microscope using a 100x or 60 x oil immersion objective and an ORCA Flash 4.0 CMOS camera (Hamamatsu). Images were acquired, processed and statistically analysed using the NIS Advanced Research software package.

### Immunoblot analysis

Saponin released schizonts were lysed in four volumes of a CoIP buffer containing: 10 mM Tris/Cl pH 7.5, 150 mM NaCl, 0.5 mM EDTA, 0.5% NP-40, 1mM PMSF, and 2x cOmplete EDTA free protease inhibitors (Roche). Lysates were incubated on ice for 10 min and then centrifuged at 12,000 x g for 20 min to remove residual solid material. Subsequently, 1 volume of 2x LDS sample buffer (Invitrogen) was added to the soluble fraction. For NBPXa processing assays, pellets were prepared as described above, and culture supernatants were first concentrated 5x in Vivaspin 500 concentrators (Sartorius) before adding LDS sample buffer. All samples were boiled for 5 min prior to separation by SDS PAGE on precast 3-8% Tris-Acetate NuPAGE gels (Invitrogen). SDS-PAGE fractionated proteins were transferred to a nitrocellulose membrane using a semidry Trans-Blot Turbo Transfer System (Bio-rad). Membranes were blocked in 10% (w/v) milk in 0.1 % PBS-Tween-20 (PBST) overnight. All primary and secondary antibodies were diluted in 1% (w/v) skimmed milk in PBST and incubated sequentially for 1 hr at room temperature followed by three washes in PBST for 10 min each. The following antibodies were used: rat α-HA (1/5000; 3F10, Sigma) and rat α-PfHSP70 (1/2000, from Ellen Knuepfer, Francis Crick Institute/RVC), followed by Goat anti-rat IgG-HRP (1/5000, Bio-Rad), and rabbit α-PkNBPXa (1/5000), followed by Goat anti-rabbit IgG-HRP (1/5000, Bio-Rad). After the final washes, membranes were incubated with Clarity Western ECL substrate (Bio-Rad) and developed using a Chemidoc imaging system (Bio-Rad).

### Live cell imaging

Purified schizonts were added to fresh human or macaque RBCs to make a 10-15% parasitaemia and 2.5% haematocrit culture. The haematocrit was subsequently adjusted to 0.25% in complete media, and 150 μl was loaded into a poly-L-lysine coated μ-Slide VI 0.4 (Ibidi). For anti-DARC assays, either 5μg/mL human anti-DARC (2C3clone, Absolute Antibody), or 5μg/mL anti-human IgG (Invitrogen) was first added to the 0.25% haematocrit culture, prior to loading samples into the channel slides. For calcium flux assays, RBCs were first incubated with 5μM fluo-4-AM (Invitrogen) in RPMI for 1h at 37°C. Cells were subsequently washed three times in RPMI, and then allowed to rest at 37°C for a further half an hour for de-esterification to occur, prior to mixing with parasites. For secretion assays performed in the absence of fresh RBCs, late stage schizonts were purified and diluted sufficiently in complete media so that egressing parasites could not contact surrounding parasitized RBCs. Loaded slides were subsequently transferred to a Nikon Ti E inverted microscope chamber, pre-warmed to 37°C. Samples were imaged using a 100x or 60 x oil immersion objective and an ORCA Flash 4.0 CMOS camera (Hamamatsu), at a rate of 1 frame/sec (100 msec exposure for each channel). Videos were acquired and processed using the NIS Advanced Research software package. Fluorescence intensities were measured using the intensity profile feature of the NIS Advanced Research software package. All other statistical analysis was carried out using Prism (version 9.0.0).

## Notes

### Competing Interest Statement

The authors have declared no competing interest.

## References

Ansari, H.R., Templeton, T.J., Subudhi, A.K., Ramaprasad, A., Tang, J., Lu, F., Naeem, R., Hashish, Y., Oguike, M.C., Benavente, E.D., Clark, T.G., Sutherland, C.J., Barnwell, J.W., Culleton, R., Cao, J., Pain, A., 2016. Genome-scale comparison of expanded gene families in Plasmodium ovale wallikeri and Plasmodium ovale curtisi with Plasmodium malariae and with other Plasmodium species. Int. J. Parasitol. 46, 685–696. https://doi.org/10.1016/j.ijpara.2016.05.009

Baker, R.P., Wijetilaka, R., Urban, S., 2006. Two Plasmodium rhomboid proteases preferentially cleave different adhesins implicated in all invasive stages of malaria. PLoS Pathog. 2, 0922–0932. https://doi.org/10.1371/journal.ppat.0020113

Beeson, J.G., Drew, D.R., Boyle, M.J., Feng, G., Fowkes, F.J.I., Richards, J.S., 2016. Merozoite surface proteins in red blood cell invasion, immunity and vaccines against malaria. FEMS Microbiol. Rev. 40, 343–372. https://doi.org/10.1093/femsre/fuw001

Chan, L.J., Dietrich, M.H., Nguitragool, W., Tham, W.H., 2020. Plasmodium vivax Reticulocyte Binding Proteins for invasion into reticulocytes. Cell. Microbiol. 22, 1–11. https://doi.org/10.1111/cmi.13110

Chen, L., Lopaticki, S., Riglar, D.T., Dekiwadia, C., Uboldi, A.D., Tham, W.H., O’Neill, M.T., Richard, D., Baum, J., Ralph, S.A., Cowman, A.F., 2011. An egf-like protein forms a complex with pfrh5 and is required for invasion of human erythrocytes by plasmodium falciparum. PLoS Pathog. 7. https://doi.org/10.1371/journal.ppat.1002199

Collins, W.E., Jeffery, G.M., 2005. Plasmodium ovale: Parasite and Disease. Clin. Microbiol. Rev. 18, 570–581. https://doi.org/10.1128/CMR.18.3.570-581.2005

Cowman, A.F., Tonkin, C.J., Tham, W.H., Duraisingh, M.T., 2017. The Molecular Basis of Erythrocyte Invasion by Malaria Parasites. Cell Host Microbe 22, 232–245. https://doi.org/10.1016/j.chom.2017.07.003

Dasgupta, S., Auth, T., Gov, N.S., Satchwell, T.J., Hanssen, E., Zuccala, E.S., Riglar, D.T., Toye, A.M., Betz, T., Baum, J., Gompper, G., 2014. Membrane-wrapping contributions to malaria parasite invasion of the human erythrocyte. Biophys. J. 107, 43–54. https://doi.org/10.1016/j.bpj.2014.05.024

Diaz, S.A., Martin, S.R., Howell, S.A., Grainger, M., Moon, R.W., Green, J.L., Holder, A.A., 2016. The binding of plasmodium falciparum adhesins and erythrocyte invasion proteins to aldolase is enhanced by phosphorylation. PLoS One 11, 1–20. https://doi.org/10.1371/journal.pone.0161850

Duffy, P.E., Patrick Gorres, J., 2020. Malaria vaccines since 2000: progress, priorities, products. npj Vaccines 5, 1–9. https://doi.org/10.1038/s41541-020-0196-3

Dvorak, J.A., Miller, L.H., Whitehouse, W.C., Shiroishi, T., 1975. Invasion of erythrocytes by malaria merozoites. Science (80-.). 187, 748–750. https://doi.org/10.1126/science.803712

Favuzza, P., de Lera Ruiz, M., Thompson, J.K., Mccauley, J.A., Olsen, D.B., Cowman, A.F., Triglia, T., Ngo, A., Steel, R.W.J., Vavrek, M., Christensen, J., Healer, J., Boyce, C., Guo, Z., Hu, M., Khan, T., Murgolo, N., Zhao, L., Penington, J.S., Reaksudsan, K., Jarman, K., Dietrich, M.H., Richardson, L., Guo, K.-Y., Lopaticki, S., Tham, W.-H., Rottmann, M., Papenfuss, T., Robbins, J.A., Boddey, J.A., Sleebs, B.E., Lè Ne, H., Sabroux, J., 2020. Dual Plasmepsin-Targeting Antimalarial Agents Disrupt Multiple Stages of the Malaria Parasite Life Cycle. Cell Host Microbe 27, 642–658. https://doi.org/10.1016/j.chom.2020.02.005

Gao, X., Gunalan, K., Yap, S.S.L., Preiser, P.R., 2013. Triggers of key calcium signals during erythrocyte invasion by Plasmodium falciparum. Nat. Commun. 4, 2862. https://doi.org/10.1038/ncomms3862

Gilson, P.R., Crabb, B.S., 2009. Morphology and kinetics of the three distinct phases of red blood cell invasion by Plasmodium falciparum merozoites. Int. J. Parasitol. 39, 91–96. https://doi.org/10.1016/j.ijpara.2008.09.007

Gruszczyk, J., Kanjee, U., Chan, L.J., Menant, S., Malleret, B., Lim, N.T.Y., Schmidt, C.Q., Mok, Y.F., Lin, K.M., Pearson, R.D., Rangel, G., Smith, B.J., Call, M.J., Weekes, M.P., Griffin, M.D.W., Murphy, J.M., Abraham, J., Sriprawat, K., Menezes, M.J., Ferreira, M.U., Russell, B., Renia, L., Duraisingh, M.T., Tham, W.H., 2018. Transferrin receptor 1 is a reticulocyte-specific receptor for Plasmodium vivax. Science (80-.). 359, 48–55. https://doi.org/10.1126/science.aan1078

Gunalan, K., Gao, X., Yap, S.S.L., Lai, S.K., Ravasio, A., Ganesan, S., Li, H.Y., Preiser, P.R., 2020. A processing product of the Plasmodium falciparum reticulocyte binding protein RH1 shows a close association with AMA1 during junction formation. Cell. Microbiol. 22. https://doi.org/10.1111/cmi.13232

Han, J.-H., Lee, S.-K., Wang, B., Muh, F., Nyunt, M.H., Na, S., Ha, K.-S., Hong, S.-H., Park, W.S., Sattabongkot, J., Tsuboi, T., Han, E.-T., 2016. Identification of a reticulocyte-specific binding domain of Plasmodium vivax reticulocyte-binding protein 1 that is homologous to the PfRh4 erythrocyte-binding domain. Sci. Rep. 6. https://doi.org/10.1038/srep26993

Haynes, J.D., Dalton, J.P., Klotz, F.W., McGinniss, M.H., Hadley, T.J., Hudson, D.E., Miller, L.H., 1988. Receptor-like specificity of a plasmodium knowlesi malarial protein that binds to duffy antigen ligands on erythrocytes. J. Exp. Med. 167, 1873–1881. https://doi.org/10.1084/jem.167.6.1873

Hillringhaus, S., Dasanna, A.K., Gompper, G., Fedosov, D.A., 2019. Importance of Erythrocyte Deformability for the Alignment of Malaria Parasite upon Invasion. Biophys. J. 117, 1202–1214. https://doi.org/10.1016/j.bpj.2019.08.027

Introini, V., Crick, A., Tiffert, T., Kotar, J., Lin, Y.C., Cicuta, P., Lew, V.L., 2018. Evidence against a Role of Elevated Intracellular Ca2+ during Plasmodium falciparum Preinvasion. Biophys. J. 114, 1695–1706. https://doi.org/10.1016/j.bpj.2018.02.023

Kazuhide, Y. N. H.M., Heledd, D., Masahito, A. C. W.S. J. T.T., Moritz, T. W. M.R., Osamu, K., 2021. Gliding motility of Plasmodium merozoites. Proc. Natl. Acad. Sci. 118, e2114442118. https://doi.org/10.1073/pnas.2114442118

Knuepfer, E., Wright, K.E., Prajapati, S.K., Rawlinson, T.A., Mohring, F., Koch, M., Lyth, O.R., Howell, S.A., Villasis, E., Snijders, A.P., Moon, R.W., Draper, S.J., Rosanas-Urgell, A., Higgins, M.K., Baum, J., Holder, A.A., 2019. Divergent roles for the RH5 complex components, CyRPA and RIPR in human-infective malaria parasites. PLoS Pathog. 15, e1007809. https://doi.org/10.1371/journal.ppat.1007809

Meyer, E.V.S., Semenya, A.A., Okenu, D.M.N., Anton, R., Bannister, L.H., Barnwell, J.W., Galinski, M.R., 2009. The reticulocyte binding-like proteins of P. knowlesi locate to the micronemes of merozoites and define two new members of this invasion ligand family. Mol. Biochem. Parasitol. 165, 111–121. https://doi.org/10.1016/j.molbiopara.2009.01.012.

The Miller, L.H., Mason, S.J., Dvorak, J.A., Mcginniss, M.H., Rothman, I.K., 1975. Erythrocyte receptors for (Plasmodium knowlesi) malaria: Duffy blood group determinants. Science (80-.). 189, 561–563. https://doi.org/10.1126/science.1145213

Mohring, F., Hart, M.N., Rawlinson, T.A., Henrici, R., Charleston, J.A., Diez Benavente, E., Patel, A., Hall, J., Almond, N., Campino, S., Clark, T.G., Sutherland, C.J., Baker, D.A., Draper, S.J., Moon, R.W., 2019. Rapid and iterative genome editing in the malaria parasite Plasmodium knowlesi provides new tools for P. vivax research. Elife 8, 1–29. https://doi.org/10.7554/elife.45829

Moon, R.W., Hall, J., Rangkuti, F., Shwen, Y., Almond, N., Mitchell, G.H., 2013. Adaptation of the genetically tractable malaria pathogen Plasmodium knowlesi to continuous culture in human erythrocytes. PNAS 110, 531–536. https://doi.org/10.1073/pnas.1216457110

Moon, R.W., Sharaf, H., Hastings, C.H., Ho, Y.S., Nair, M.B., Rchiad, Z., Knuepfer, E., Ramaprasad, A., Mohring, F., Amir, A., Yusuf, N.A., Hall, J., Almond, N., Lau, Y.L., Pain, A., Blackman, M.J., Holder, A.A., 2016. Normocyte-binding protein required for human erythrocyte invasion by the zoonotic malaria parasite Plasmodium knowlesi. Proc. Natl. Acad. Sci. U. S. A. 113, 7231–7236. https://doi.org/10.1073/pnas.1522469113

Ngernna, S., Chim-Ong, A., Roobsoong, W., Sattabongkot, J., Cui, L., Nguitragool, W., 2019. Efficient synchronization of Plasmodium knowlesi in vitro cultures using guanidine hydrochloride. Malar. J. 18, 1–7. https://doi.org/10.1186/s12936-019-2783-1

Nichols, M.E., Rubinstein, P., Barnwell, J., Rodriguez de Cordoba, S., Rosenfield, R.E., 1987. A new human Duffy blood group specificity defined by a murine monoclonal antibody. Immunogenetics and association with susceptibility to Plasmodium vivax. J. Exp. Med. 166, 776–785. https://doi.org/10.1084/jem.166.3.776

O’Donnell, R.A., Hackett, F., Howell, S.A., Treeck, M., Struck, N., Krnajski, Z., Withers-Martinez, C., Gilberger, T.W., Blackman, M.J., 2006. Intramembrane proteolysis mediates shedding of a key adhesin during erythrocyte invasion by the malaria parasite. J. Cell Biol. 174, 1023–1033. https://doi.org/10.1083/jcb.200604136

Pal-Bhowmick, I., Andersen, J., Srinivasan, P., Narum, D.L., Bosch, J., Miller, L.H., 2012. Binding of Aldolase and Glyceraldehyde-3-Phosphate Dehydrogenase to the Cytoplasmic Tails of Plasmodium falciparum Merozoite Duffy Binding-Like and Reticulocyte Homology Ligands. MBio 3, 1–8. https://doi.org/10.1128/mBio.00292-12.Editor

Perrin, A.J., Collins, C.R., Russell, M.R.G., Collinson, L.M., Baker, D.A., Blackman, M.J., 2018. The actinomyosin motor drives malaria parasite red blood cell invasion but not egress. MBio 9, 1–13. https://doi.org/10.1128/mBio.00905-18

Rawlinson, T.A., Barber, N.M., Mohring, F., Cho, J.S., Kosaisavee, V., Gérard, S.F., Alanine, D.G.W., Labbé, G.M., Elias, S.C., Silk, S.E., Quinkert, D., Jin, J., Marshall, J.M., Payne, R.O., Minassian, A.M., Russell, B., Rénia, L., Nosten, F.H., Moon, R.W., Higgins, M.K., Draper, S.J., 2019. Structural basis for inhibition of Plasmodium vivax invasion by a broadly neutralizing vaccine-induced human antibody. Nat. Microbiol. 4, 1497–1507. https://doi.org/10.1038/s41564-019-0462-1

Reddy, K.S., Amlabu, E., Pandey, A.K., Mitra, P., Chauhan, V.S., Gaur, D., 2015. Multiprotein complex between the GPI-anchored CyRPA with PfRH5 and PfRipr is crucial for Plasmodium falciparum erythrocyte invasion. Proc. Natl. Acad. Sci. U. S. A. 112, 1179–1184. https://doi.org/10.1073/pnas.1415466112

Riglar, D.T., Richard, D., Wilson, D.W., Boyle, M.J., Dekiwadia, C., Turnbull, L., Angrisano, F., Marapana, D.S., Rogers, K.L., Whitchurch, C.B., Beeson, J.G., Cowman, A.F., Ralph, S.A., Baum, J., 2011. Super-resolution dissection of coordinated events during malaria parasite invasion of the human erythrocyte. Cell Host Microbe 9, 9–20. https://doi.org/10.1016/j.chom.2010.12.003

Singh, A.P., Ozwara, H., Kocken, C.H.M., Puri, S.K., Thomas, A.W., Chitnis, C.E., 2005. Targeted deletion of Plasmodium knowlesi Duffy binding protein confirms its role in junction formation during invasion. Mol. Microbiol. 55, 1925–1934. https://doi.org/10.1111/j.1365-2958.2005.04523.x

Singh, S., Alam, M.M., Pal-Bhowmick, I., Brzostowski, J.A., Chitnis, C.E., 2010. Distinct external signals trigger sequential release of apical organelles during erythrocyte invasion by malaria parasites. PLoS Pathog. 6. https://doi.org/10.1371/journal.ppat.1000746

Smolarek, D., Hattab, C., Hassanzadeh-Ghassabeh, G., Cochet, S., Gutiérrez, C., de Brevern, A.G., Udomsangpetch, R., Picot, J., Grodecka, M., Wasniowska, K., Muyldermans, S., Colin, Y., Le Van Kim, C., Czerwinski, M., Bertrand, O., 2010. A recombinant dromedary antibody fragment (VHH or nanobody) directed against human Duffy antigen receptor for chemokines. Cell. Mol. Life Sci. 67, 3371–3387. https://doi.org/10.1007/s00018-010-0387-6

Tham, W.H., Healer, J., Cowman, A.F., 2012. Erythrocyte and reticulocyte binding-like proteins of Plasmodium falciparum. Trends Parasitol. 28, 23–30. https://doi.org/10.1016/j.pt.2011.10.002

Tham, W.H., Lim, N.T.Y., Weiss, G.E., Lopaticki, S., Ansell, B.R.E., Bird, M., Lucet, I., Dorin-Semblat, D., Doerig, C., Gilson, P.R., Crabb, B.S., Cowman, A.F., 2015. Plasmodium falciparum Adhesins Play an Essential Role in Signalling and Activation of Invasion into Human Erythrocytes. PLoS Pathog. 11, 1–22. https://doi.org/10.1371/journal.ppat.1005343

Triglia, T., Tham, W.H., Hodder, A., Cowman, A.F., 2009a. Reticulocyte binding protein homologues are key adhesins during erythrocyte invasion by Plasmodium falciparum. Cell. Microbiol. 11, 1671–1687. https://doi.org/10.1111/j.1462-5822.2009.01358.x

Triglia, T., Tham, W.H., Hodder, A., Cowman, A.F., 2009b. Reticulocyte binding protein homologues are key adhesins during erythrocyte invasion by Plasmodium falciparum. Cell. Microbiol. 11, 1671–1687. https://doi.org/10.1111/j.1462-5822.2009.01358.x

Volz, J.C., Yap, A., Sisquella, X., Thompson, J.K., Lim, N.T.Y., Whitehead, L.W., Chen, L., Lampe, M., Tham, W.H., Wilson, D., Nebl, T., Marapana, D., Triglia, T., Wong, W., Rogers, K.L., Cowman, A.F., 2016. Essential Role of the PfRh5/PfRipr/CyRPA Complex during Plasmodium falciparum Invasion of Erythrocytes. Cell Host Microbe 20, 60–71. https://doi.org/10.1016/j.chom.2016.06.004

Wanaguru, M., Crosnier, C., Johnson, S., Rayner, J.C., Wright, G.J., 2013. Biochemical analysis of the plasmodium falciparum erythrocyte-binding antigen-175 (EBA175)-glycophorin-A interaction; Implications for vaccine design. J. Biol. Chem. 288, 32106–32117. https://doi.org/10.1074/jbc.M113.484840

Weiss, G.E., Gilson, P.R., Taechalertpaisarn, T., Tham, W.H., de Jong, N.W.M., Harvey, K.L., Fowkes, F.J.I., Barlow, P.N., Rayner, J.C., Wright, G.J., Cowman, A.F., Crabb, B.S., 2015. Revealing the Sequence and Resulting Cellular Morphology of Receptor-Ligand Interactions during Plasmodium falciparum Invasion of Erythrocytes. PLoS Pathog. 11, 1–25. https://doi.org/10.1371/journal.ppat.1004670

Wertheimer, S.P., Barnwell, J.W., 1989. Plasmodium vivax interaction with the human Duffy blood group glycoprotein: Identification of a parasite receptor-like protein. Exp. Parasitol. 69, 340–350. https://doi.org/https://doi.org/10.1016/0014-4894(89)90083-0

Williams, A.R., Douglas, A.D., Miura, K., Illingworth, J.J., Choudhary, P., Murungi, L.M., Furze, J.M., Diouf, A., Miotto, O., Crosnier, C., Wright, G.J., Kwiatkowski, D.P., Fairhurst, R.M., Long, C.A., Draper, S.J., 2012. Enhancing Blockade of Plasmodium falciparum Erythrocyte Invasion: Assessing Combinations of Antibodies against PfRH5 and Other Merozoite Antigens. PLoS Pathog. 8. https://doi.org/10.1371/journal.ppat.1002991

