## Supplementary Figures for "Sequential roles for red blood cell binding proteins enable phased commitment to invasion for malaria parasites"

**Supplementary Data**

**List of Supplementary Videos**

- Video S1: *P. knowlesi* invasion of human RBC
- Video S2: *P. knowlesi* merozoites gliding on human RBCs with and without deformation
- Video S3: *P. knowlesi* merozoite invading a Fluo-4-AM pre-loaded human RBC example 1
- Video S4: *P. knowlesi* merozoites invading macaque RBCs
- Video S5: DMSO vs. Rap treated NBPXa cKO merozoites interacting with human RBCs
- Video S6: *P. knowlesi* merozoites interacting with human RBCs with anti-DARC (Fy6)
- Video S7: NBPXa-mNG secretion without merozoites interacting with RBCs
- Video S8: DBPα-mNG secretion without merozoites interacting with RBCs
- Video S9: Invasion of human RBC by NBPXa-mNG tagged merozoite
- Video S10: Invasion of human RBC by DBPα-mNG tagged merozoite

**Supplementary Figures**

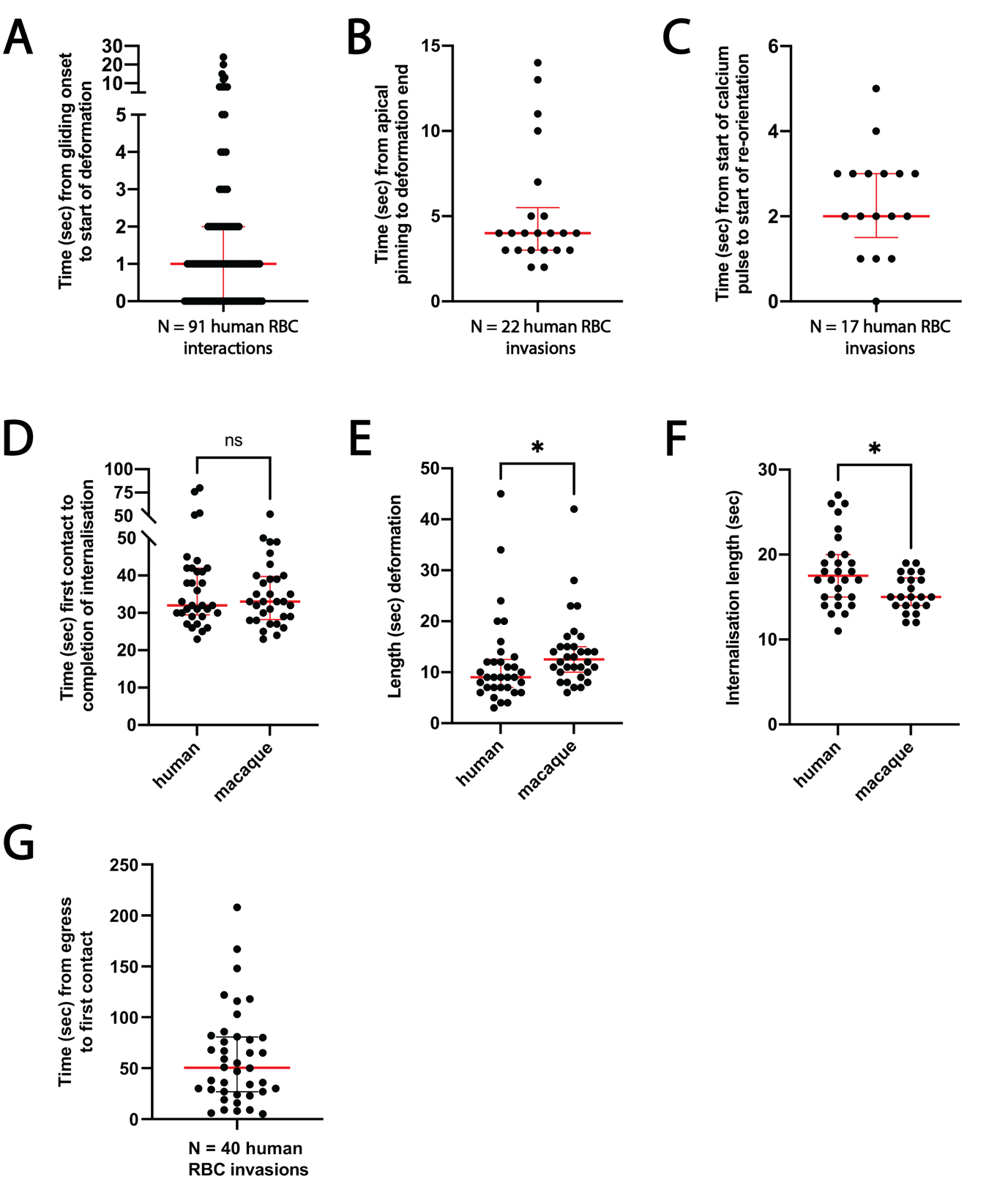

**S1. Analysis of *P. knowlesi* interactions with human and macaque RBCs (A)** Merozoites spend a median of 1 second gliding on human RBCs (N = 91 interactions; IQR = 0-2 sec). **(B)** Merozoites cease forward movement across human RBC surfaces a median 4 sec (IQR 3-5.5 sec) before deformation subsides. N = data from 22 invasions. **(C)** Merozoites re-orientate on human RBC surfaces a median 2 sec (IQR = 2-5 sec) after a calcium pulse is observed. N = data from 17 invasions. **(D)** There is no significant difference between the length of time from first contact to completion of internalisation for human (median = 32 sec; IQR = 29.5-42 sec) vs. macaque (median = 33 sec; IQR = 28-40 sec) invasions. ‘Ns’ indicates p = 0.722, when analysed using a Mann-Whitney U-test. **(E)** *P. knowlesi* merozoites spend significantly longer deforming macaque RBCs (median = 12.5 sec; IQR = 10-15 sec; N = 32 invasions) vs. human (median = 9 sec; IQR = 7-12.5 sec; N = 33 invasions) prior to internalisation. * indicates p = 0.016, when analysed using a Mann-Whitney U-test. **(F)** *P. knowlesi* merozoites spend significantly longer actively invading human RBCs (mean = 18.1 sec; SD = 4.3 sec; N = 28 invasions) vs. macaque (mean = 15.5 sec; SD = 2.2 sec; N = 22 invasions). * indicates p = 0.010, when analysed using unpaired t-test. **(G)** Merozoites typically contact the human RBC they will invade within a median 50 sec post egress (IQR = 27-80.75 sec).

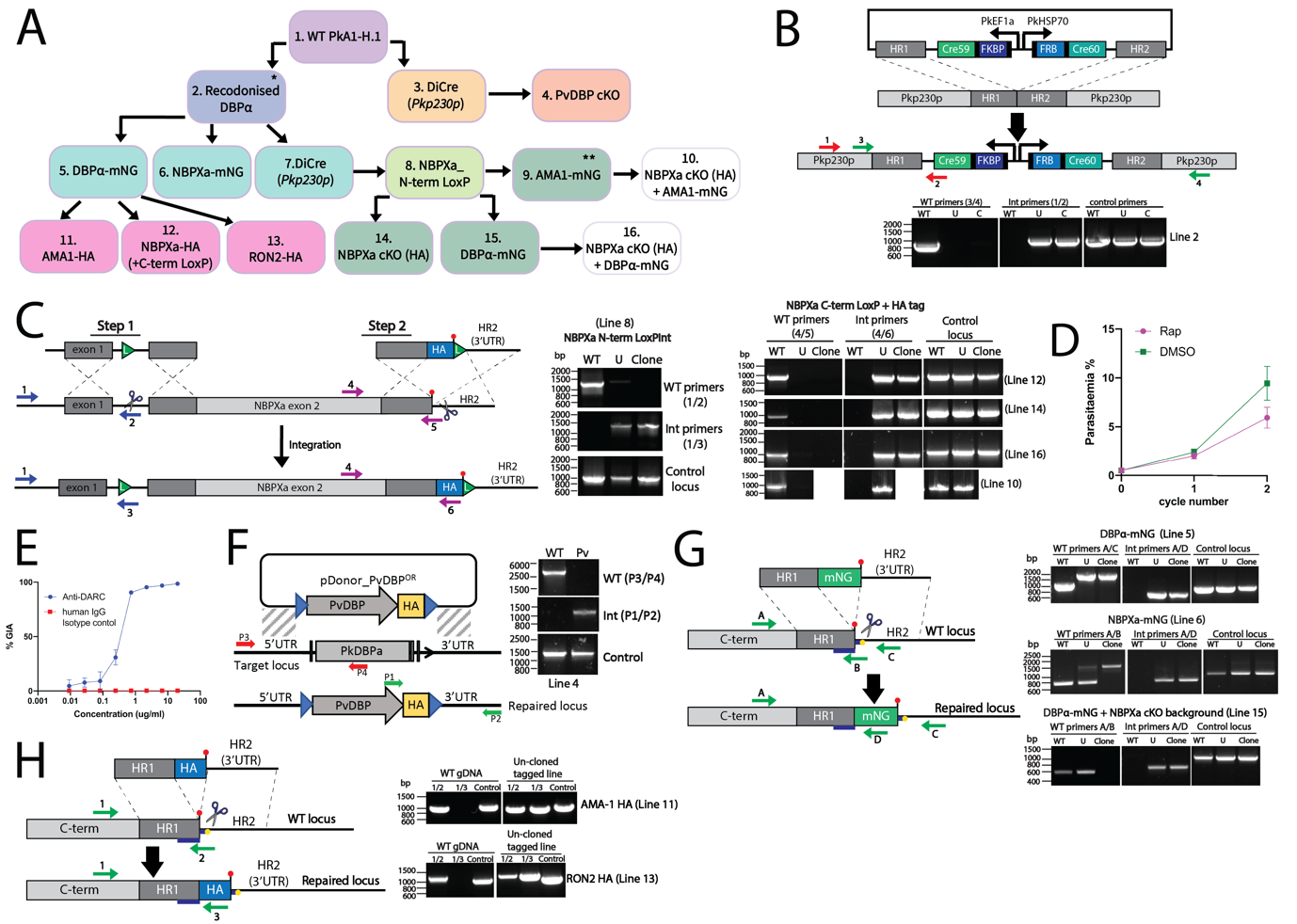

**S2. Generation and genotyping of *P. knowlesi* transgenic lines (A)** Flow chart of transgenic lines generated/used in this study. *indicates line described in Mohring et al., 2019. **indicates line described in Yahata and Hart et al., 2021. **(B)** Generation of Dimerisable Cre Recombinase (DiCre) expressing backgrounds (lines 2 and 3). Transgenic lines were screened with primers FM49 and FM50 (positions 3 and 4 in schematic) to detect WT parasites (expected band size = 859 bp) and primers FM90 and FM32 (positions 1 and 2 in schematic) to detect integrated parasites (expected band size = 891 bp). **(C)** Generation of conditional NBPXa cKO/HA tagged lines. Lines were generated with two steps (see schematic). Step 1 = integrating a LoxP site into the endogenous NBPXa intron. WT parasites detected with primers MH959 and MH1143 (positions 1 and 2 in schematic; expecting 1261 bp band). Integrated parasites detected with primers MH959 and MH1172 (positions 1 and 3 in schematic; expected band size = 1284 bp). Step 2 = integrating a LoxP sequence + single HA tag at the C-term of NBPXa. Primers MH1129 and MH1128 were used to detect WT parasites (positions 4 and 5 in schematic; expected band size = 1004bp), and primers MH1129 and MH563 were used to detect integrated parasites (positions 4 and 6 in schematic; expected band size = 997 bp). Lines 10, 14, and 16 were full conditional KO lines (contained both steps 1 and 2). Line 12 was transfected into line 5 background (eg. wasn’t a cKO line – just had C-term LoxP + HA tag). **(D)** Treating ‘WT’ Dicre parasites (Line 8) with rapamycin induces a slight, but non-significant growth defect (p = 0.122, when comparing final parasitaemias by unpaired t-test; N = results from 5 independent assays). **(E)** Representative growth inhibition assay (GIA) results showing dose dependent inhibition of *P. knowlesi* invasion of human RBCs with increasing concentrations of human anti-DARC, Fy6 antibody (2C3 clone, Absolute antibody). EC50 = 0.3 ug/ml (range = 0.23-0.36 ug/ml, from 4 independent experiments). **(F)** Generating a PvDBP cKO line. WT PkDBPα was swapped with a floxed PvDBP sequence (schematic). Primers FM1973 + FM188 (positions 3 and 4 in schematic) were used to detect WT parasites (expected size = 2773 bp). Primers FM1897 and FM1041 were used to detect integration of PvDBP construct in cloned parasites ‘Pv’ (expected size = 980 bp). **(G)** Schematic showing the insertion of an mNG tag prior to the stop codon (red circle) of *Pk* invasion genes. Blue lines indicate the position of the Cas9 guide sequences. For DBPa mNG, line 5, primers FM421 and FM225 were used to detect WT parasites (primers in positions A/C; expected band size = 1788bp if integrated or 1046 bp if WT). Primers FM421 and MH291 were used to detect integrated parasites (primers in positions A/D; expected band size = 691 bp). For DBPa mNG, line 15, primers FM421 and MH1130 were used to detect WT parasites (primers in positions A/B; expected band size = 564 bp). Primers FM421 and MH291 were used to detect integrated parasites, as above. For NBPXa mNG, line 6, primers MH835 and 1128 were used to detect WT parasites (primers in positions A/B; expected band size = 863 if WT & 1565 bp if integrated). Primers MH835 and MH291 were used to detect integration (primers in positions A/D; expected size = 973 bp). For all PCRs, ‘U’ = uncloned parasites. **(H)** Schematic showing the insertion of a HA tag at the C-terminus of PkAMA-1 and PkRON2. For AMA-1, primers MH892 and MH904 were used to detect WT parasites (primers in positions 1 and 2 in schematic. Expected band size = 984 bp). Primers MH892 and MH563 used to detect integrated parasites (primers in positions 1/3; expected band size = 996 bp). For RON2, primers MH890 and MH909 were used to detect WT parasites (primers in positions 1 and 2 in schematic. Expected band size = 1209 bp). Primers MH890 and MH563 used to detect integrated parasites (primers in positions 1/3; expected band size = 1338 bp).

**Supplementary Tables**

**Supplementary Table 1.** Primers, DNA templates, and expected PCR product sizes for generating transfection constructs (white cells) and screening transgenic *P. knowlesi* parasites (green cells).

| **PCR Product** | **Fwd primer** | **Rev primer** | **product size (bp)** | **Template(s)** |
| --- | --- | --- | --- | --- |
| mNeonGreen tag | | | | |
| mNG PCR product | MH297 | MH295 | 836 | mNeonGreen plasmid (Yahata and Hart et al., 2019) |
| PkNBPXa-mNG Donor | | | | |
| HR1 | MH227 | MH229 | 514 | WT PkA1H1 gDNA |
| HR2 | MH306 | MH231 | 553 | WT PkA1H1 gDNA |
| HR1-mNG fusion | MH227 | MH296 | 1222 | HR1 + mNG PCR products |
| HR1-mNG-HR2 fusion | MH298 | MH299 | 1566 | HR1-mNG + HR2 PCR products |
| ‘WT’ locus | MH835 | MH1128 | 863 | gDNA from transfection |
| Integrated locus | MH835 | MH291 | 973 | gDNA from transfection |
| PkDBPα-mNG Donor | | | | |
| HR1 | MH221 | MH223 | 524 | Recodonised PkDBPα gDNA (Mohring et al., 2019) |
| HR2 | MH224 | MH225 | 521 | Recodonised PkDBPα gDNA  (Mohring et al., 2019) |
| HR1-mNG fusion | MH221 | MH296 | 1232 | HR1 + mNG PCR products |
| HR1-mNG-HR2 fusion | MH222 | MH226 | 1637 | HR1-mNG + HR2 PCR products |
| WT locus | FM421 | MH225 | 1046 | gDNA from transfection |
| WT locus | FM421 | MH1130 | 564 | gDNA from transfection |
| Integrated locus | FM421 | MH291 | 691 | gDNA from transfection |
| PkRON2-HA Donor | | | | |
| HR1 | MH847 | MH849 | 1092 | WT PkA1H1 gDNA |
| HR2 | MH850 | MH851 | 1007 | WT PkA1H1 gDNA |
| HR1-HR2 fusion | MH848 | MH852 | 1874 | HR1 + HR2 PCR products |
| WT locus | MH890 | MH909 | 1209 | gDNA from transfection |
| Integrated locus | MH890 | MH563 | 1338 | gDNA from transfection |
| PkAMA-1-HA Donor | | | | |
| HR1 | MH841 | MH843 | 872 | WT PkA1H1 gDNA |
| HR2 | MH844 | MH845 | 880 | WT PkA1H1 gDNA |
| HR1-HR2 fusion | MH842 | MH846 | 1512 | HR1 + HR2 PCR products |
| WT locus | MH892 | MH904 | 984 | gDNA from transfection |
| Integrated locus | MH892 | MH563 | 996 | gDNA from transfection |
| PkNBPXa N-term LoxP Donor | | | | |
| HR1 | MH953 | MH967 | 1037 | WT PkA1H1 gDNA |
| HR2 | MH966 | MH957 | 974 | WT PkA1H1 gDNA |
| HR1-HR2 fusion | MH954 | MH958 | 1651 | HR1 + HR2 PCR products |
| WT locus | MH959 | MH1143 | 1261 | gDNA from transfection |
| Integrated locus | MH959 | MH1172 | 1284 | gDNA from transfection |
| PkNBPXa C-termLoxP + HA tag Donor | | | | |
| HR1 | MH835 | MH238 | 858 | WT PkA1H1 gDNA |
| HR1 extended | MH835 | MH837 | 895 | HR1 PCR product |
| HR2 | MH838 | MH839 | 852 | WT PkA1H1 gDNA |
| HR1ext-HR2 fusion | MH975 | MH1023 | 1507 | HR1e + HR2 PCR products |
| WT locus | MH1129 | MH1128 | 1004 | gDNA from transfection |
| Integrated locus | MH1129 | MH563 | 997 | gDNA from transfection |
| PvDBP cKO + HA tag Donor | | | | |
| C-term LoxP | FM0167 | FM1893 | 618 | pDonor_PvDBP^OR^ (Rawlinson et al., 2019) |
| C-term LoxP extended 1 | FM1895 | FM1894 | 574 | C-term LoxP |
| C-term LoxP extended 2 | FM1897 | FM1896 | 247 | C-term LoxP extended 1 |
| N-term LoxP | FM0141 | MH1788 | 686 | pDonor_PvDBP^OR^ (Rawlinson et al., 2019) |
| N-term LoxP extended | FM1899 | FM1898 | 652 | N-term LoxP |
| WT locus | FM1973 | FM188 | 2773 | gDNA from transfection |
| Integrated locus | FM1897 | FM1041 | 980 | gDNA from transfection |
| DiCre Cassette Donor | | | | |
| Cre59 | FM268 | FM267 | 536 | pBS_DC_hsp86-Bip5 (Collins et al., 2013) |
| Cre60 | FM265 | FM266 | 1220 | pBS_DC_hsp86-Bip5 (Collins et al., 2013) |
| EF1α 5’UTR | FM269 | FM270 | 646 | PkconGFPp230p (Moon et al., 2013) |
| PbDT 3’UTR | FM271 | FM272 | 864 | PkconGFPp230p (Moon et al., 2013) |
| PfHRP2 3’UTR | FM273 | FM274 | 587 | PkconGFPp230p (Moon et al., 2013) |
| PkHSP70 5’UTR | FM275 | FM276 | 1345 | PkconGFPp230p (Moon et al., 2013) |
| p230p HR1 | FM047 | FM053 | 417 | WT PkA1H1 gDNA |
| P230p HR2 | FM089 | FM006 | 431 | WT PkA1H1 gDNA |
| WT locus | FM49 | FM50 | 859 | gDNA from transfection |
| Integrated locus | FM90 | FM32 | 891 | gDNA from transfection |
| NBPXa C-term Guide plasmid | | | | |
| Piece 1 | MH400 | MH236 | 442 | pL_11HF |
| Piece 2 | MH235 | MH295 | 704 | pL_11HF |
| Full length insert | FM401 | FM285 | 662 | Piece 1 + Piece 2 PCR products |

**Supplementary Table 2.** Sequences of primers used in this study.

| **Primer** | **Sequence** |
| --- | --- |
| MH297 | GGATCCGGTGGAGGCAGCGG |
| MH295 | CGACAGGTTTCCCGACTGGAAAG |
| MH227 | GAAAATAAATCTATTAGAGGAAGAGGAAGTTAAGC |
| MH306 | ACAGATGTTATGGGAATGGATGAATTGTATAAATAATGAGTAAGGGGGATAGAAGTTTATACAAAAAG |
| MH298 | CAGTTACTAGCAATTTGAATGAGCAGT |
| MH229 | ACCTCCGCTGCCTCCACCGGATCCTATATATTCGTTACTTTCGTCAAAGGTAACTTCA |
| MH231 | TTCGTTTTTCAAAATATGTCTTTTAGGCACC |
| MH296 | TTATTTATACAATTCATCCATTCCCATAACATCTGT |
| MH299 | AGTTTCTCTGATTATCTAATTAATTGATATAAATTCCCATC |
| MH221 | ACAACCGGAAACTGAACCGG |
| MH224 | GGAATGGATGAATTGTATAAATAATGATGCTACTTGGGTAAGTAAGGAGA |
| MH222 | TTTATCCGCGGAGAAGATCATCCTGATGAGCGAAG |
| MH223 | ACCTCCGCTGCCTCCACCGGATCCGCTGTAGTCAAGGGGGGTGAACTG |
| MH225 | CGGGGCTAATTTGTCCGTGTA |
| MH226 | TTTATGAATTCGACAAATTAATGGCACATTTTTCTCTTTTGG |
| MH847 | TTCAGTGTAGCTAGAAACCCGGG |
| MH848 | GCTGATCATTGTTCTGCCTTGTG |
| MH851 | TGCAAAATCCTCTTGTTTTCCCC |
| MH852 | GGTTCAAAATAGGATAAGTCAAAACGGC |
| MH841 | CAACCGAATCGTTTTCGAAGCTAG |
| MH842 | TGATTAAAAGTGCCTTTCTTCCAGTGG |
| MH845 | TATGGATTATATTAAAGTAAGCGTATACGCCATG |
| MH846 | TAGAGTGCTCATATAAGGCACGCAC |
| MH850 | TACCCATACGATGTTCCAGATTACGCTTAGGGTGGTGACTTGTCCATCTC |
| MH849 | AGCGTAATCTGGAACATCGTATGGGTACATCTGTATGCGGGCGTACGA |
| MH844 | TACCCATACGATGTTCCAGATTACGCTTGAGTGGGGAAGCAACGTATTATTCG |
| MH843 | AGCGTAATCTGGAACATCGTATGGGTAGTAGTAAGGCTTCTCCATCAGAACAG |
| MH953 | CTTGTGGTGCTCTTTTAGGTAAAATTACC |
| MH954 | TTATCCGCGGGAAGCTGTTTGAGGTAATCGTGTG |
| MH957 | CAGCCGAATTGCACTCCTTAG |
| MH958 | TTATGCGGCCGCCCTGTTCGTTACCTTCTGTCTGTG |
| MH966 | CTTCGTATAGCATACATTATACGAAGTTATATGCTAAACACCCCTTTATGTTAAACC |
| MH967 | TCGTATAATGTATGCTATACGAAGTTATAGGATGTATAATAATGCCATATAGATCAGCC |
| MH835 | GGCAAGATAGAAGCATATATCGAAAACATTTC |
| MH838 | ATAGCATACATTATACGAAGTTATGTAAGGGGGATAGAAGTTTATACAAAAAGAAAATC |
| MH975 | TTATCCGCGGGAATCCTATAATGAAGAGGCAAGGAAAAAATTAC |
| MH238 | AGCGTAATCTGGAACATCGTATGGGTATATATATTCGTTACTTTCGTCAAAGGTAACTTCA |
| MH837 | ATAACTTCGTATAATGTATGCTATACGAAGTTATTCAAGCGTAATCTGGAACATCGTATG |
| MH839 | CGTATTTCCCCATAAAGATGAATGCG |
| MH1023 | TTATGCGGCCGCGAAAATTATGAAAACGCCATGTTTAAATTTGC |
| FM0167 | CGAGCGGAGAAAGCACATGAACAA |
| FM1893 | TACGAAGTTATTCAAGCGTAATCTGGAACATCGTATGGGTAGCTGTAGTCCAGGGGGGTG |
| FM1895 | CCTGTCCAACGGAAAGCTGGAC |
| FM1894 | GCGGCCGCATAACTTCGTATAATGTATGCTATACGAAGTTATTCAAGCGTAATCTGGAAC |
| FM1897 | CTGCTGCTGATCGCCAGCAGA |
| FM1896 | CTCCTTACTTACCCAAGTAGCAGCGGCCGCATAACTTCGTATAATGTAT |
| FM0141 | GTTTTCCCAGTCACGAC |
| MH1788 | CTTCGTATAATGTATGCTATACGAAGTTATTGTATTTAAAAATAAGTGTATACG |
| FM1899 | CACTATAGGGCGAATTGGCGGAAG |
| FM1898 | TCTTGCCCTTCATACTAGTATAACTTCGTATAATGTATGCTATACGAAGTTATTG |
| FM268 | ATATGGTACCTCAGTTCAGCTTGCACCAGGCA |
| FM267 | ATATCTCGAGATGGCCCCTAAGAAGAAGAGAAAGG |
| FM265 | ATATCTCGAGATGGCCCCTAAGAAGAAGAGAAAGG |
| FM266 | ATATGGTACCTCAGTCCCCATCCTCCAGCAG |
| FM047 | ATATCCGCGGCAGAAGCCGGGTTAGCAGCAC |
| FM053 | ACTAGTTGGAACCCACTTCGGGTGGTC |
| FM89 | ATATGCTGAGCATGAACCTCTGAGCGAAGAGGAGC |
| FM006 | TTTTACCGTTCCATGGGGCGCGCCACATGGCACTCCTTATTATCCTTCGTTAG |
| FM269 | AGGATATGCGGCCGCTAAGTAACCCTTGCATATGCCCCT |
| FM270 | AGGATATACTAGTATAGGTACCATACTCGAGTTTCGAATAAAATTAAATTGAAAAAAAGGTAAGTACGGG |
| FM271 | ATATGGTACCATATGGCAGCTTAATGTTCGTTTTTCTTATTTATATATTT |
| FM272 | ATATGCTCAGCCTACCCTGAAGAAGAAAAGTCCGATG |
| FM273 | ATATGGTACCATTTAATAATAGATTAAAAATATTATAAAAATAAAAACATAAACACAGAA |
| FM274 | ATATACTAGTTAGATTTAATAAATATGTTCTTATATATAATGAGAAATAAATATTTAACA |
| FM275 | GGATATGCGGCCGCATGCAATATACCCATTTTGAATACACCCCA |
| FM276 | GGATATGCTCAGCATAGGTACCATACTCGAGTTTTACGGGGATCTGCAAGGGG |
| MH400 | TATGCCTAAGATCGTCTCCCCTTTGATT |
| MH236 | TTCTAGCTCTAAAACTACTCATATATATTCGTTACAATAATATACTGTAA |
| MH235 | TTACAGTATATTATTGTAACGAATATATATGAGTAGTTTTAGAGCTAGAA |
| MH295 | CGACAGGTTTCCCGACTGGAAAG |
| FM401 | CATTGTTCCCCCCTTTGTTTTGCAAG |
| FM285 | TACTTATGCGTATACAAAGCCTTCTTCAC |
| MH835 | GGCAAGATAGAAGCATATATCGAAAACATTTC |
| MH291 | TTGGATTTCCTGTTCCTTGTCCAACC |
| FM421 | TTCATGCGGAACAGCAACAG |
| MH225 | CGGGGCTAATTTGTCCGTGTA |
| MH1130 | ACCCAAGTAGCATCAGCTGTAG |
| MH890 | TGC TAG AAA AAG TGG CAG AGT TAC ATA AG |
| MH909 | AGT CAC CAC CCT ACA TCT GTA TGC |
| MH563 | CGTAATCTGGAACATCGTATGGG |
| MH892 | GTA ATG ATT GGG AAA ACA AGT GCC C |
| MH904 | GCT TCC CCA CTC AGT AGT AAG G |
| MH959 | CCCCTTTCCCAGGAACAAATTG |
| MH1143 | CATAAAGGGGTGTTTAGCATAGGATG |
| MH1172 | GTTTAGCATATAACTTCGTATAATGTATGCTATACGAAG |
| MH1129 | GAAGGCATATACGAAATATGGAAAAGAGC |
| MH1128 | CTTTTTGTATAAACTTCTATCCCCCTTACTCATATATATTC |
| FM1973 | CCATGTACACGATTTGTGTACTTATAGAATC |
| FM188 | AGGAGCACCTGATTGAGAACCTGGA |
| FM1897 | CTGCTGCTGATCGCCAGCAGA |
| FM1041 | GTAGGGAACATTTCTTTCTGCGG |
| FM49 | GAAGATTCCGCAAAGCTTTGTCGGTTA |
| FM50 | ACGCTATGGAAGCAGTTGTCTGGAT |
| FM90 | TGGGAAATACAGGAAATAACGGTGTTATGT |
| FM32 | CCTAATCATGTAAATCTTAAATTTTTCTTTTTAAACATATG |
